# Dopaminergic reward signals shape foraging decisions through learned value expectations

**DOI:** 10.64898/2026.10.21.760765

**Authors:** Lyle Kingsbury, Hugo May, Naoshige Uchida

## Abstract

A fundamental function of the nervous system is to direct animals towards life-sustaining resources and enable adaptive strategies for acquiring them. Foraging, the challenge of securing these resources from the environment, requires making decisions that balance local gains against costs, opening a window on value-based decision making in ecological context. Yet despite the centrality of foraging in life, the algorithms and neural mechanisms underlying foraging decisions are not well understood. We studied mice making stay-or-leave patch foraging decisions for water across poor- and rich-resource environments with dynamic local reward outcomes. We found that mice integrate immediate rewards, local reward history, and reward (i.e. value) expectations to guide foraging decisions, favoring greater time investment in rich-environment patches. Computational models revealed influence of a time-varying bias which mirrored the shape of a learned value function. Consistent with a reinforcement learning mechanism, dopamine in the ventral striatum signaled a temporal-difference reward prediction error, which could be used as a teaching signal to construct value functions, and calibrated optogenetic manipulation of dopamine to create fictive reward signals recapitulated the value-related decision bias. Taken together, our results shed new light on the neurocomputational basis of foraging, revealing a novel role for value expectations in dynamic environments. Further, they establish generalized function of biological reinforcement learning in ecological context, demonstrating a specific role for the canonical dopamine reward circuit in shaping temporally resolved value expectations to guide behavior and cognition. The unification of this framework with foraging opens a new avenue to elucidate principles of how animals learn and dynamically adjust cognitive strategies for real-world ecological problems.

## Introduction

A fundamental function of the nervous system is to direct animals towards life-sustaining resources (“rewards” such as food and water) and enable adaptive strategies for acquiring them. Foraging, the challenge of finding and harvesting resources from the environment, requires tracking reward information and making decisions that balance resource gains against costs of opportunity, energy, and threat.^1,2^ Ubiquitous in nature and across species, foraging behaviors provide an ecological window onto value-based decision processes in the brain, which are central to life in the animal kingdom and in modern human societies.^3–5^

Foraging problems have been studied extensively at the behavioral level, shedding light on how animals make decisions in natural settings. In *patch foraging*, a forager navigates its environment to find resource locations (patches), and as resources deplete locally, must decide when to leave in search of alternatives (**Figure 1A**). This canonical stay-or-leave decision problem has been analyzed theoretically,^1,2,6,7^ and recent studies have begun to reveal its neural basis. In humans and primates, correlates of foraging variables have been identified in the frontal cortex;^8–10^ and in rodents, neural dynamics can compute patch leaving decisions by integrating time and reward gains to a decision boundary,^11–13^ akin to drift-diffusion processes for perceptual choices.^14,15^ However, major explanatory gaps persist between theory and mechanistic understanding. While models often assume foragers have knowledge of reward statistics, such as values and reward rates, it remains unclear how these are learned and represented at the neural level,^9,10,16^ and while such value-related variables can flexibly adjust foraging strategies,^17–20^ the cellular and circuit mechanisms underlying this remain unknown.

**Figure 1.**
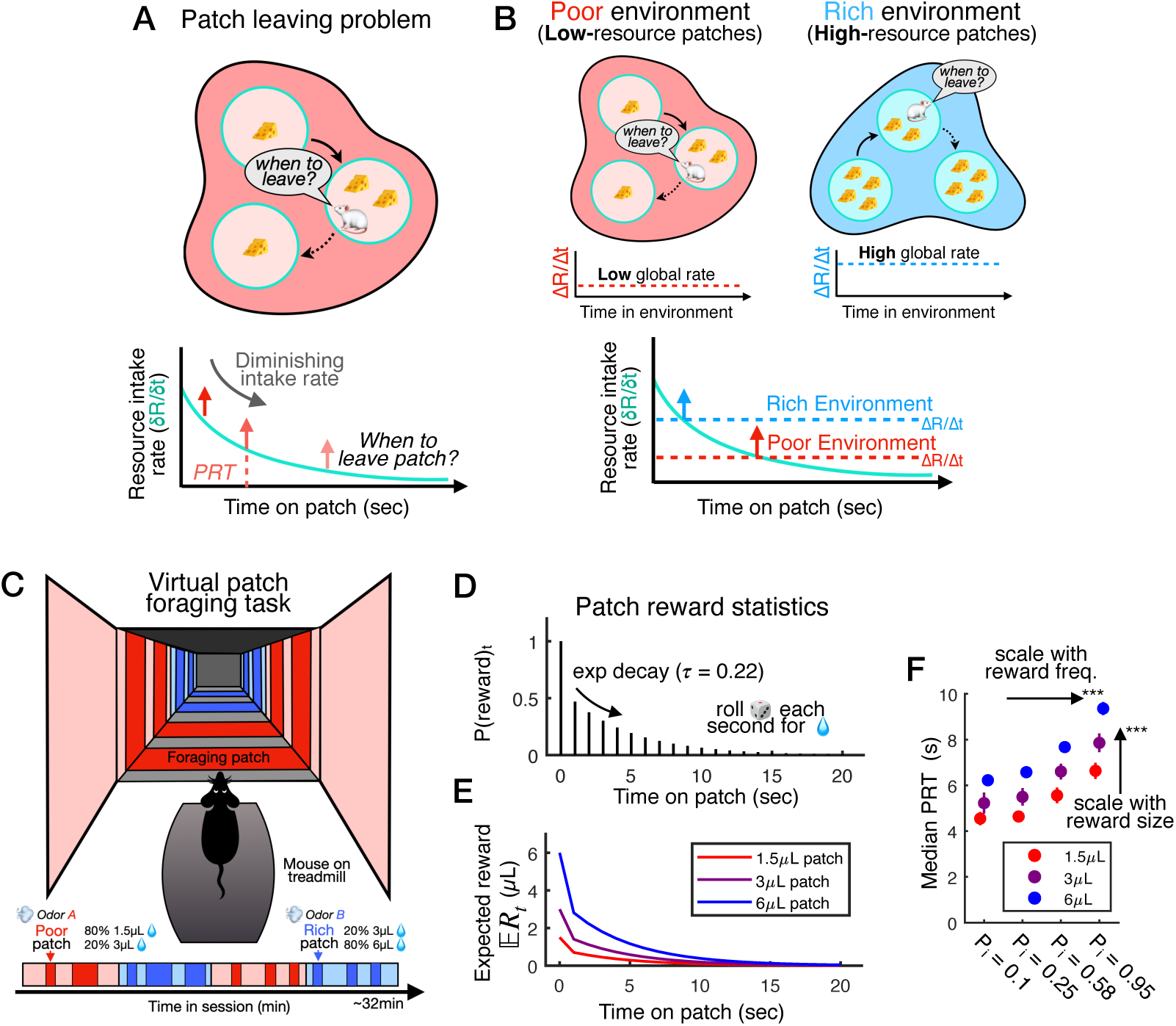
Patch foraging decisions across poor and rich resource environments. **A)** Cartoon of the canonical patch leaving problem, in which a forager navigates their environment to find localized resource patches, and as the resource (reward, R) intake rate (δ**R/**δ**t**) diminishes within a patch, they must decide when to leave in search of alternatives. The time invested before leaving at a given patch is the patch residence time (PRT, seconds). **B)** Animals forage across poor- and rich-resource environments with low and high global reward rates (**ΔR/Δt**). To maximize long-term reward gains, animals should adapt their leaving decisions and investment times to long-term reward statistics, as they represent different opportunity costs for continuing to harvest a patch (also see **Figure S1** about the *Marginal Value Theorem*, MVT). **C)** A virtual patch foraging task for mice with a poor environment (red, with 1.5µL and 3µL reward patches) and a rich environment (blue, with 3µL and 6µL reward patches). Poor and rich environments, and poor and rich patches within, are distinguished by visual and odor cues. Mice forage across both in alternating blocks in daily sessions. **D)** When mice stop at a patch, they receive one water reward at t = 0s, and then can receive additional rewards each second with decaying probability (exponential decay from t = 1s, τ = 0.22s). **E)** Patch quality (small, medium, large) is determined by the size of water drops (1.5µL, 3µL, 6µL), producing distinct functions of decaying expected reward E**R_t_** for each type. All rewards within a patch are the same size. **F)** Empirical PRTs measuring time invested on patches with increasing reward frequency (initial reward probability at t = 1s, P_i_ on x-axis) and reward size (color), showing that mice leave later on patches with higher returns (median ± s.d. for each size/ probability patch type, multiple linear regression, *P* = 4.82e-19 [reward size], *P* = 1.28e-24 [reward probability], *n* = 22879 patch trials pooled across sessions).

An influential framework posits that animals construct internal representations of expected future reward (i.e. value) through an iterative reinforcement learning (RL) process.^21,22^ In the mammalian brain, mesolimbic dopamine (DA) is thought to drive RL by broadcasting a temporal-difference (TD) reward prediction error (RPE) teaching signal,^23–28^ positioning DA circuitry as an attractive candidate substrate underlying value learning for foraging. Yet although much is known about how DA-mediated RL can promote behavioral reinforcement^29–31^ and forge associations between stimuli and reward outcomes,^32–35^ this TD-RL model has been formulated and refined largely in the context of simple conditioning paradigms, and it less clear whether or how the core RL algorithm and circuit implementation^34^ operate in more complex, real-world settings.^36–40^ Furthermore, while DA plays a critical role in value learning, it is also important for movement, motivation, and cognition,^35,41–44^ and as debate continues, the specific computational form and function of DA signals in these diverse aspects of natural behavior remain to be clarified.^45–48^

To understand the computational and neural basis of value-based foraging decisions, we developed a virtual foraging paradigm to study mice making patch leaving decisions in resource-poor vs. rich environments with dynamic reward rates. We found that mice exhibit a bias toward greater time investment in rich-environment patches, indicating a value effect on foraging decisions directionally opposite that predicted by optimal foraging theory. Computational models suggest that this bias reflects influence of a learned value function that encodes time-varying reward expectations separately from immediate reward outcomes and local reward history. Hypothesizing a critical role for mesolimbic reward circuity, we found that DA in the ventral striatum (VS) reflects this value function through a TD-RPE signal. Calibrated stimulation of midbrain DA neurons to produce fictive reward signals recapitulated the learned, value-based decision bias without affecting movement or online decision making. Together, these results demonstrate operation of DA-mediated TD-RL for self-paced foraging, establishing generality of the theoretical framework for natural behavior in ecological context. Furthermore, they clarify a specific role of phasic DA reward signals in driving value learning for foraging, linking this canonical function of the TD-RL circuit to the flexible control of a fundamental cognitive process.

## Main

### Patch foraging decisions are sensitive to local and global reward rates

Foraging is a ubiquitous natural behavior that requires efficiently balancing immediate resource gains against costs, such the opportunity to forage elsewhere. In the canonical patch foraging problem (**Figure 1A**), a forager navigates its environment to find localized resource patches (e.g. yielding water/food rewards), and as intake diminishes with consumption, must decide when to leave in search of alternatives. Fundamentally, patch leaving decisions involve a comparison of reward information across different scales: *immediate local gains* from staying at the current patch must be compared against *potential future gains* resulting from leaving the patch. Many species, including rodents and humans,^10,11,49^ solve patch foraging problems efficiently and exhibit sensitivity to local reward outcomes, investing more time on patches with higher yield. Additionally, because the opportunity cost of staying is related to potential future gains, leaving decisions should also be sensitive to the overall reward rate of the environment (the global reward rate), leading to different decision strategies in rich vs. poor-resource environments (**Figure 1B**).

The framework of optimal foraging theory offers a normative way to conceptualize this: for example, the Marginal Value Theorem (MVT) states that the optimal patch leaving time occurs when the local (marginal) reward rate equals the long-term environment (global) reward rate (**Figure S1A**).^1,2^ Thus, environment richness is predicted to generally reduce patch residence times (PRT, how long a forager stays at a given patch, **Figure 1B, S1A**), as has been observed in several foraging studies.^8,10,50–53^ We confirmed this prediction by simulating patch leaving decisions across different environments under an MVT-based decision policy (**Figure S1B-C**).

Yet despite the utility of idealized models for guiding intuitions, empirical behavior is inconsistent across experiments and species, and it remains unclear how animals forage in naturalistic environments with stochastic or dynamic reward outcomes.^54–57^ Further, while optimality theories assume that foragers know (or can estimate) the relevant reward statistics,^1,2^ the mechanisms by which these are learned, represented, and used to shape foraging decisions are not well understood.^10,16^

### A virtual foraging task with dynamic local and global reward statistics

To elucidate the computational and neural mechanisms underlying value-based patch leaving decisions, we designed an experimental paradigm in which mice foraged for water rewards across resource-poor and rich environments with stochastic local reward outcomes (**Figure 1C**). Head-fixed mice navigate a virtual corridor and stop at visual/odor-cued patches, and once stopped, they receive water rewards each second with decaying probability and must decide when to leave (**Figure 1D**, top).^11,12^ While all patches yield rewards with the same decaying probability, the poor environment contains small (1.5µL water, 80%) and medium (3µL, 20%) reward patches, while the rich environment contains mostly large reward patches (6µL, 80%, **Figure 1C-E**). Comparison of behavior across reward-matched 3µL patches in the poor vs. rich environment thus provides insight into the effect of the environment on leaving decisions. In each session, mice foraged for 30-40 minutes across both environments in alternating blocks. Consistent with our previous reports,^11,12^ mice exhibited stable variability in average PRTs and patch engagement (**Figure S2A-D**), and they positively scaled PRTs with reward size and frequency, indicating strong sensitivity to immediate reward outcomes on individual patch trials (**Figure 1F**).

To test whether mice distinguish poor vs. rich environments, we examined motivated behavior in the task by comparing patch engagement rates (the fraction of patches mice visited) and the speed with which mice moved between patches (**Figure S2E**). Mice showed significantly higher engagement in the rich environment (**Figure S2F**) and moved more quickly between rich patches (**Figure S2G**). This shows that mice are more motivated to forage for water rewards in the rich environment, indicating an implicit understanding of environment-specific reward statistics and flexible adaptation of behavior.

### Mice spend more time harvesting reward-matched patches in the rich environment

We next studied patch leaving decisions by comparing PRTs across trial types, focusing on reward-matched 3µL patches in poor vs. rich environments. Surprisingly, and contrary to the theoretical prediction based on the MVT (**Figure S1**),^1,2^ mice invested significantly more time on 3µL patches in the rich environment (**Figure 2A**, late days). Although the general effect of immediate rewards (**Figure 1F**) was present early on, this rich environment PRT bias emerged over time (**Figure 2A-C**), suggesting that it may be related to learned reward expectations shaped by experience.

**Figure 2.**
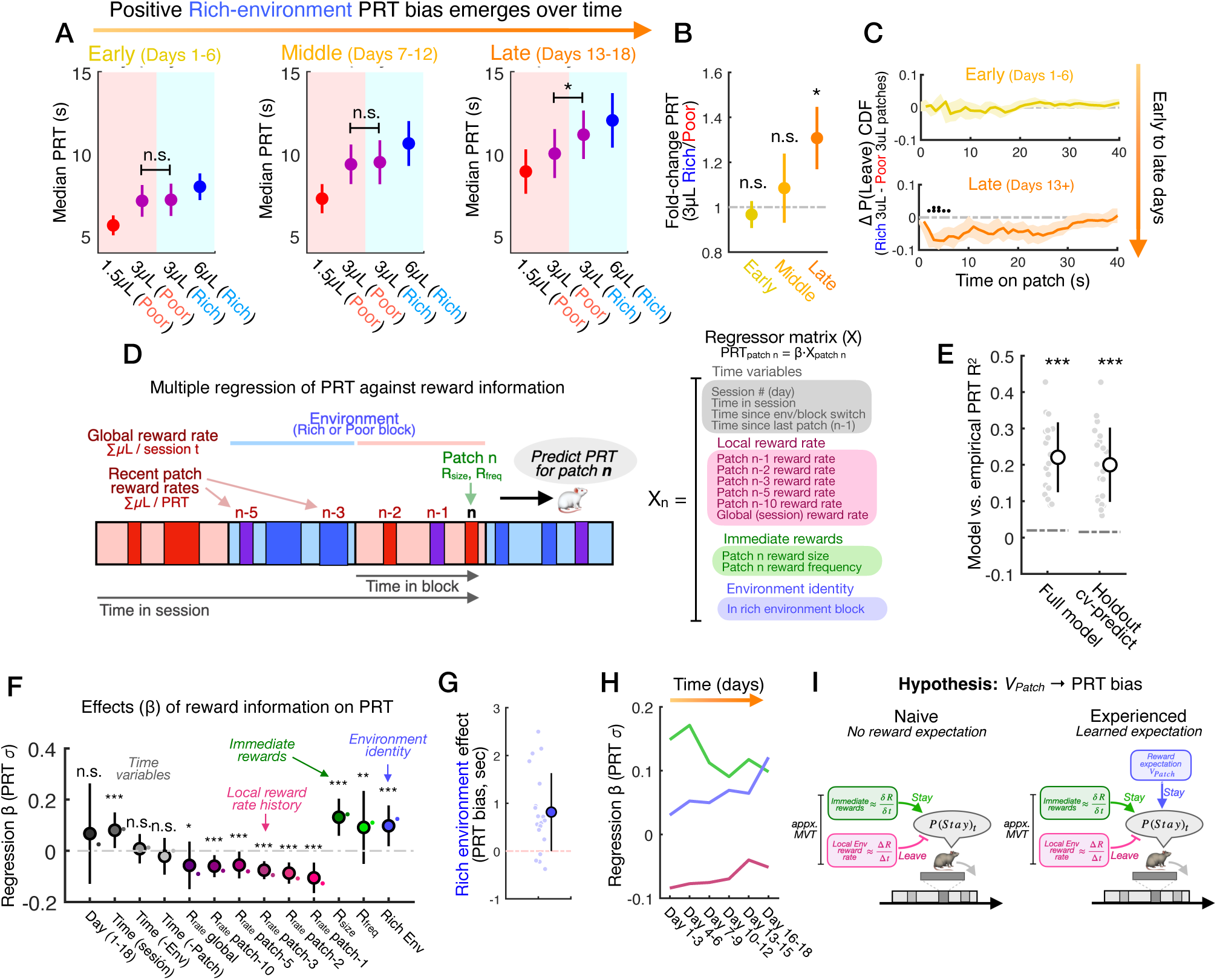
Mice exhibit positively biased patch residence times in the rich environment. **A)** Patch residence times (PRT) for small (1.5µL), medium (3µL), and large (6µL) patches across poor and rich environments on early (days 1-6), middle (days 7-12), and late (days 13-18) sessions. On late days, mice exhibit significantly longer PRTs on reward-matched 3µL patches in the rich vs. poor environment (rich-environment PRT bias, two sample t-tests, 3µL poor vs. rich, *P* = 0.91 [early], *P* = 0.864 [middle], *P* = 0.012 [late], *n* = 20 mice). **B)** Rich PRT bias across early, middle, and late days, quantified as the ratio of average PRTs (for each mouse) on 3µL patches in the rich vs. poor environment (one sample t-tests, *P* = 0.59 [early], *P* = 0.587 [middle], *P* = 0.038 [late], *n* = 20 mice). **C)** Rich-environment PRT bias across sessions shown as the difference in P(Leave) CDFs for 3µL patches in the rich vs. poor environment. **D)** Left, illustration of regression analysis to predict single-trial PRTs as a function of time- and reward-related task variables; right, regressor set used for the model, including time variables (grey), local reward rate history (pink), immediate current-patch rewards (green), and cued patch reward information as environment identity (rich vs. poor, blue). **E)** Performance (R^2^) of regression model to explain and predict trial PRTs (one model for each mouse, *P* = 8.86e-5 [full model vs. shuffle], *P* = 8.86e-5, [holdout predictions vs. shuffle], SR tests, *n* = 20 mice). **F)** Coefficient β fits (average across models for each mouse) for regressors schematized in (**D**) to model trial PRTs (mean ± s.d., one sample t-tests, *P* = 1.77e-7 [1-back reward rate], *P* = 1.31e-7 [current patch reward size], *P* = 0.0098 [current patch reward frequency], *P* = 2.8e-5 [rich environment block], *n* = 20 mice, see also **Table S1**). **G)** The average effect size in seconds of the positive rich-environment PRT bias captured by the regression analysis (mean ± s.d., µ = 0.816s, *P* = 2.59e-4, *n* = 20 mice). **H)** Evolution of β coefficients over time (3-day bins), averaged within regressor sets representing different reward information (local reward rate, immediate rewards, environment bias). The rich environment bias increases over time while the effect of immediate rewards (frequency and reward size on the current patch) decreases over time. **I)** The direction and time course of the rich environment PRT bias raise the hypothesis that in experienced mice, a learned value expectation **V_Patch_** biases decisions away from patch leaving (longer patch time investment), in addition to the effects of immediate rewards and the local environment reward rate present in the naive state.

To understand how reward statistics and expectations jointly influence patch leaving decisions, we used multiple regression to quantitatively analyze the correlations between reward variables and PRT behavior. For each patch trial, we computed regressors that capture reward statistics across different timescales (i.e. immediate patch rewards, local reward history from recent patches, and environment identity, which could cue reward expectations) and used linear regression to model trial PRTs (**Figure 2D**). The model captured and predicted PRTs significantly above chance levels (**Figure 2E**), so we next analyzed the coefficients (**β**) fit to reward regressors to understand their relationship with (and potential influence on) leaving decisions.

We first confirmed that immediate rewards (current patch reward size and frequency) positively affect PRT (**Figure 2F**). By contrast, regressors capturing the local reward rate history on recent patches carried a decaying negative influence, suggesting that mice are also sensitive to local reward rates and, within individual sessions, leave patches earlier when the local reward rate is higher.^56^ Together, this suggests that within an environment, mice implement a computation for leaving decisions that weighs immediate patch gains against the local reward rate, consistent with the logic of the MVT (**Figure S1**). However, even when accounting for this, the rich environment regressor was significantly positive (**Figure 2F**), indicating that the observed rich patch PRT bias is not driven by environment reward rate *per se*, but rather by cued environment identity.

The effect size for this regressor was around 1 second (**Figure 2G**), consistent with the average difference in PRTs across 3µL patches (**Figure 2A**) and indicating that both analyses capture the same bias effect. Indeed, when we constructed the model for different session bins across time, we observed an evolution in coefficients consistent with the average PRT trend (**Figure 2H**): while the effect of immediate and local reward history was already present early (pink and green, days 1-3), the rich bias (blue) was initially small and increased progressively with task experience.

The positive direction and temporal evolution of the rich environment PRT bias thus raise the hypothesis that a learned expectation of reward outcomes (i.e. a value expectation) might independently influence patch leaving decisions (**Figure 2I**). This value effect could bias mice toward greater investment in patches they expect more rewards from, irrespective of actual reward outcomes, reducing their sensitivity to the inherent variability of unpredictable, stochastic rewards.

### A learned patch value function might influence patch leaving decisions

To explore this idea, we turned to the framework of reinforcement learning (RL).^21,22^ RL theory formalizes reward expectation as a learned representation of value ***V*** for a given state of the world ***s*** (or temporal state ***t***), with a value function ***V_s_*** representing values over states (**Figure 3A**). State value is defined as the expected sum of rewards yielded by that state and states that follow, discounted into the future (by ***γ***). For each state, ***V_s_*** thus represents its total reward expectation under a diminishing time horizon. In principle, value functions can be learned from experienced reward outcomes using RL algorithms (**Figure 3A**). At the biological level, evidence suggests that the mammalian brain constructs value expectations using an RL process, which is thought to depend critically on teaching signals from midbrain dopamine (DA) neurons.^23,24,27,28,45^

**Figure 3.**
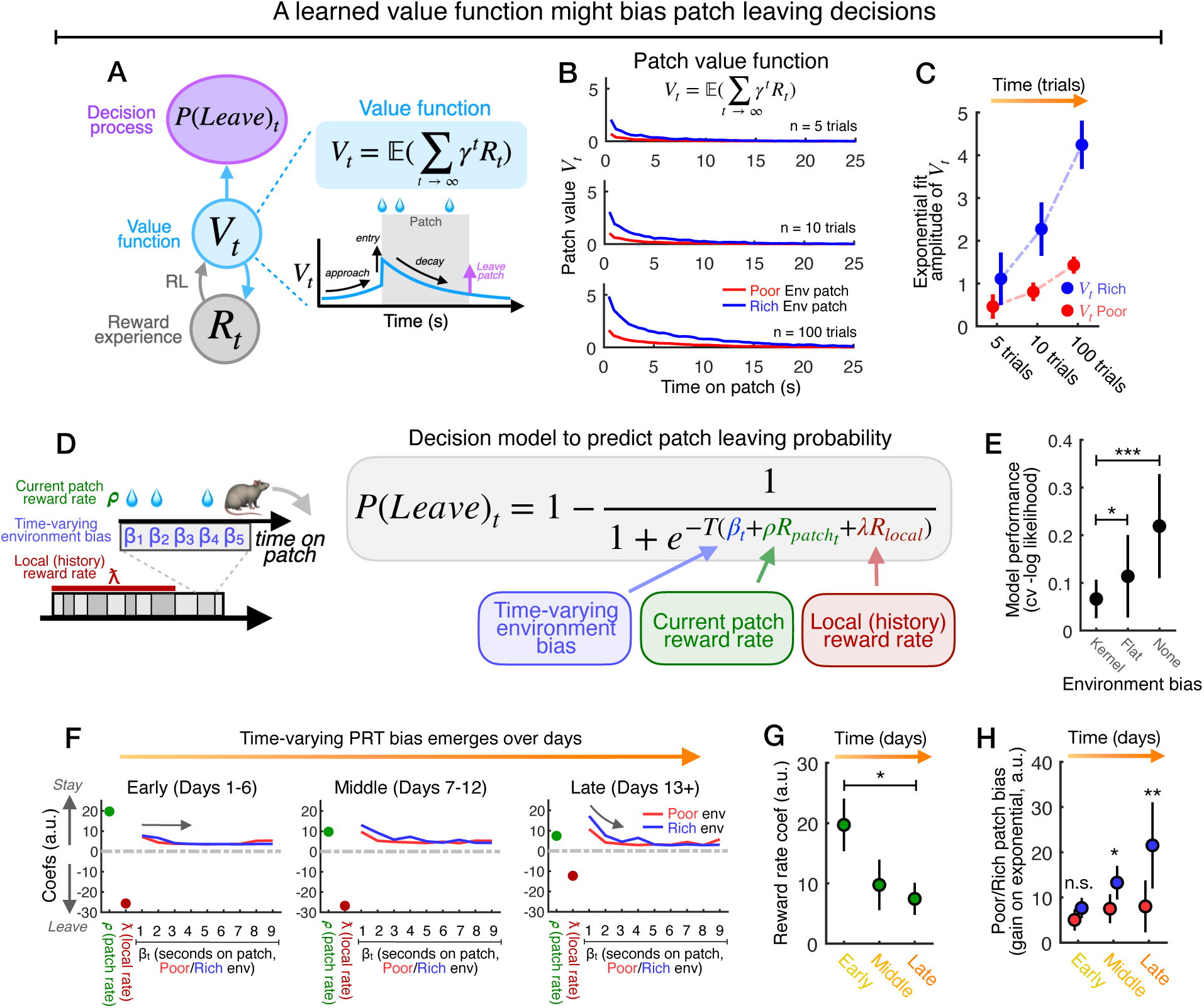
Bias from a learned value function explains the patterns of decision behavior. **A)** Hypothesized model in which a value function **V_t_**biases mice toward longer PRTs by inhibiting the leaving decision **P(Leave)_t_**. The value function **V_t_** encoding expected future (discounted) reward at each time point can be constructed from reward experience **R_t_** through a reinforcement learning (RL) algorithm. B) Simulated evolution (after 5, 10, and 100 trials) of a patch value function **V_t_** by application of the temporal difference (TD) RL algorithm to empirical reward outcomes experienced by mice in the foraging task (**Methods**). C) The value functions for poor and rich patches develop and diverge over time, with the rich patch function converging to a higher value (amplitude α of an exponential fit to the value function, mean ± s.d., *n* = 100 simulations). D) Decision model to predict time-varying patch leaving probability **P(Leave)_t_** using reward-related variables over the course of individual patch trials. The effect of poor vs. rich environment identity (the linear bias in the PRT regression, Figure 2F) is expanded as a time-varying effect captured by bias parameters **β_t_** tiling the patch in 1-second bins, in addition to the current patch reward rate (weighted by ρ) and local reward rate on recent patches (weighted by ƛ). E) Model comparison measuring performance to predict **P(Leave)_t_** in held-out data (cross-validated -log likelihood). The decision model with the time-varying bias (Kernel) significantly out-performs reduced models with either a constant bias (Flat) or no bias (mean ± s.d., two sample t-tests, *P* = 0.044 [Kernel vs. Flat], *P* = 5.31e-6 [Kernel vs. None], *n* = 20 mice). F) Parameter fits for the decision model across early (days 1-6), middle (days 7-12), and late (days 13-18) sessions. The positive and negative weights on patch reward rate (green) and local reward rate history (pink), respectively, follow the same pattern as in the PRT regression (Figure 2F). The expanded PRT bias, captured by **β_t_**, follows the emergence of a shape like the patch value function (**B-C**): flat on early days (left) and an exponential decay pattern on late days (right), with a higher amplitude on rich (blue) vs. poor (red) environment patches. G) Evolution of the weight parameter ρ for current patch reward rate across days, showing progressively diminished weight of immediate reward outcomes on patch leaving decisions (SR test, *P* = 0.016, *n* = 20 mice, see also Figure 2H). H) Evolution of the environment-specific time-varying bias captured by **β_t_**, and quantified as the amplitude **ɑ** of an exponential fit to **β_t_** coefficients for poor and rich patches for each bin (early, middle, late). The pattern of the bias on leaving decisions tracks the emergence of an exponential shape and divergence across poor and rich patches, matching the development of poor and rich patch value functions (shuffle tests for difference in poor vs. rich **ɑ**, *P* = 0.052 [early], *P* = 0.002 [middle], *P* = 0.001 [late], *n* = 20 mice, 1000 shuffles, see also **B-C** and **Methods**).

We hypothesized that a value function ***V_t_*** representing time-varying reward expectation might underlie the observed PRT bias on rich patches (**Figure 3A**), which emerged over time and was independent of immediate rewards. Because patches in our task yield rewards with decaying probability (***τ*** = 0.22, **Figure 1E**), such a value function would have specific features reflecting this structure: increasing value on patch approach, a rapid increase on patch entry, and a decay with time, reflecting diminishing reward expectation (**Figure 3A**). To understand how a value function might emerge through RL, we performed a simulation using the temporal difference (TD) learning algorithm applied to the empirical reward outcomes experienced by mice in our task (**Figure 3B**).^21,22^ Value functions for poor and rich environment patches were initially flat and developed an exponential shape over time, with progressively higher amplitude for rich patches reflecting greater reward expectations (**Figure 3C**).

### A time-varying bias driven by a learned value function explains decision behavior

The hypothesis that a value function drives the rich PRT bias carries several specific predictions: **1)** the PRT bias should be stronger on rich vs. poor patches; **2)** it should decay with time on the patch, rather than be constant, reflecting the shape of the value function; and **3)** these two features should be absent initially and emerge only with learning, tracking an RL process.

To test these, we formulated a decision model for patch leaving (**Figure 3D**) which constructs a time-varying leaving probability ***P***(***Leave***)***_t_*** as a threshold over a linear weighted combination of three factors: the current patch reward rate ***R_patch_***, the local reward rate on recent patches ***R_local_***, and environment-specific, time-varying biases tiling seconds on the patch ***β_t_***. In this model, the set of bias weights ***β_t_*** define a flexible kernel that captures the influence of poor vs. rich patch identity on leaving decisions, allowing us to measure the shape of PRT biases in each environment and test the specific predictions of the value function hypothesis.

We first compared performance of different versions of the model to predict leaving probabilities. The model with a time-varying bias performed better than models with constant (i.e. flat) bias or no bias (**Figure 3E**), indicating that poor vs. rich identity explains non-redundant variance in decision behavior, consistent with the regression (**Figure 2D-F**). Analyzing the full bias kernel captured by ***β_t_***, we observed a striking correspondence with features of the patch value function: while on early days the bias was nearly flat and similar on poor vs. rich patches (**Figure 3F**, left), a time-varying structure emerged on later days, approximating an exponential decay with higher amplitude on rich patches (**Figure 3F**, right). Indeed, while influence of the current reward rate ***R_patch_*** again decreased over days, the bias effect increased, with progressively higher amplitude on rich vs. poor patches (**Figure 3H**)

These results support a picture in which the rich PRT bias reflects influence of a learned patch value function on leaving decisions, effectively increasing time investment on patches with high expected reward, independent of immediate reward outcomes. However, although the behavioral pattern is consistent with this, whether the brain represents this value function, and the specific neural mechanisms driving its construction, remain unknown.

### Ventral striatal dopamine signals the RL error signal and reflects the value function

To address these questions, we next studied dopamine (DA) release in the ventral striatum (VS), which is known to play an important role in reward learning. In this model of biological RL, value expectations are learned through a DA teaching signal that takes the form of a temporal-difference reward prediction error (TD-RPE, **Figure 4A**).^22,34,38,45^ The TD-RPE ***δ_t_*** reflects deviation from a value expectation ***V_t_*** as a subtractive contrast between actual and expected reward: ***δ_t_*** = ***R_t_***_+**1**_ + ***γV_t_***_+**1**_ − ***V_t_***. Algorithmically, ***δ_t_*** iteratively updates ***V_t_***, producing a better estimate of the value function with more reward experience. At the biological level, TD-RPE signals are hypothesized to drive neural plasticity and modify representations of expected value in recipient neuron populations,^58,59^ such as in the VS.^34^ If this TD-RL system is indeed used to learn value functions for foraging, then the VS DA signal should exhibit specific properties consistent with TD-RPE, including transient responses to unexpected rewards and changes in expected value, as well as negative modulation by a time-varying value estimate (i.e. the value function ***V_t_***).^24–27^

**Figure 4.**
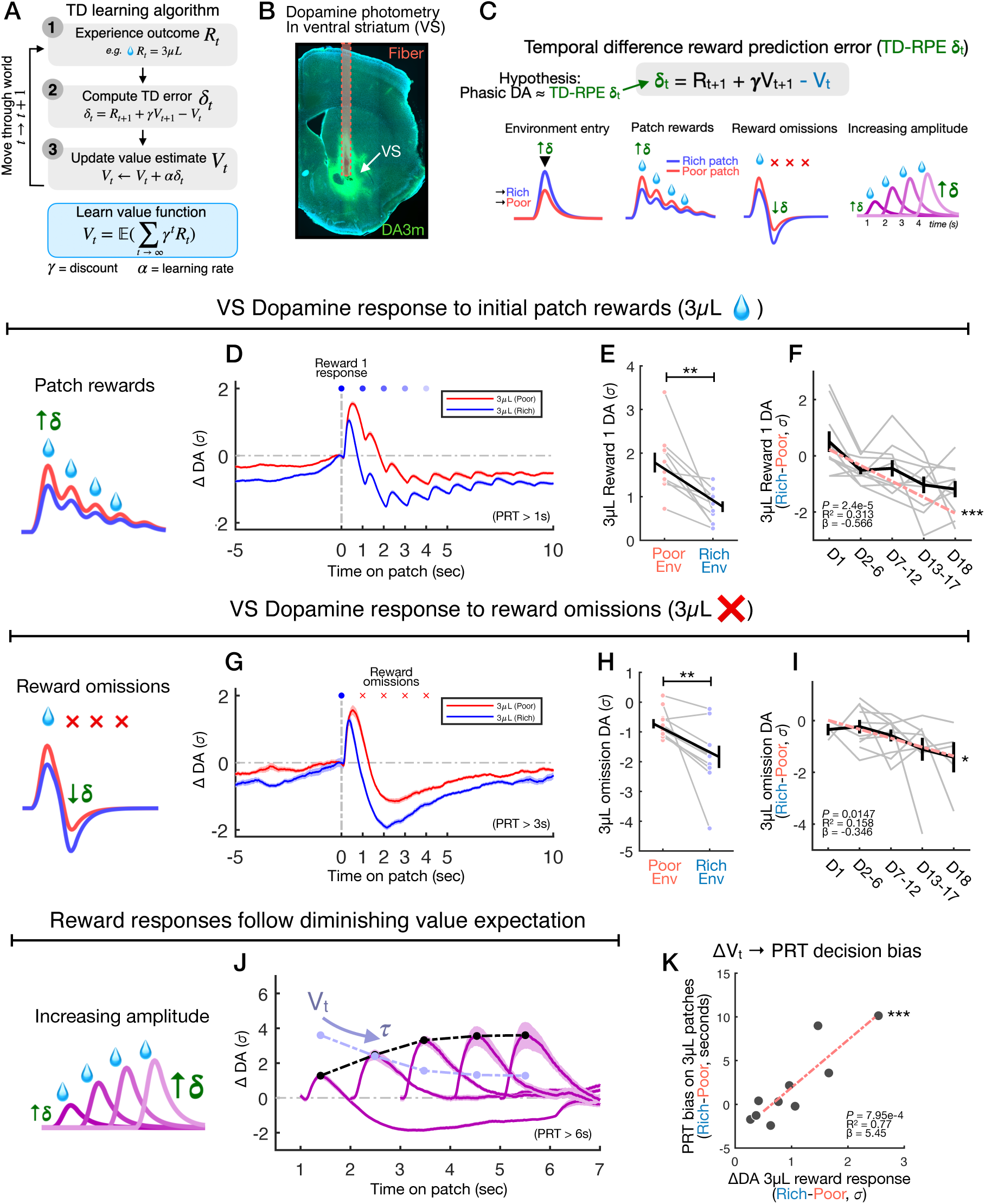
Ventral striatal dopamine reflects value through a temporal difference reward prediction error. **A)** Schematic of the temporal difference (TD) algorithm for value learning. Reward outcomes **R_t_** are experienced as an agent moves through states of the world, and the TD reward prediction error (RPE) *δ***_t_** is computed as the sum of current reward **R_t_** and the discounted value of the next state γ**V_t+1_** minus the current value estimate **V_t._** *δ***_t_** is used to update the value estimate **V_t_** to minimize future surprise. The phasic component of the dopamine signal is theorized to encode a TD-RPE teaching signal (i.e. *δ***_t_**) to drive value learning in the brain. B) Experimental strategy to record dopamine (DA) release in the ventral striatum (VS) during naturalistic foraging, and histological section from an example mouse showing expression of the DA fluorescence sensor GrabDA3m (green) and placement of an optic fiber to collect the photometric VS DA signal. C) Cartoon illustrating predicted patterns of VS DA around events in the foraging task under the hypothesis that phasic DA responses correspond to an approximate TD-RPE (δ**_t_**). The TD-RPE δ**_t_** should be positive upon entry to foraging environments (left); *δ***_t_** responses to water rewards and omissions should be modulated by poor vs. rich context (middle); and *δ***_t_** reward responses should increase with time, reflecting the diminishing value expectation **V_t_** (right, see also Figure 3B). D) Trial-average of the VS DA signal aligned to the trial-initial water reward on medium-reward 3µL patches in the poor (red) or rich (blue) environment (late sessions, days 13-18). The trial-initial reward response is expressed in z-scores as a change from the pre-trial baseline (ΔDA, σ, mean ± SEM, *n* = 362 poor, *n* = 405 rich, see also **Figure S3E**). E) Comparison of the difference in 3µL reward responses across poor vs. rich patches (paired comparison across mice, mean ± s.d., measured at 0.5s, SR test, *P* = 0.002, *n* = 10 mice). F) The difference between poor vs. rich patch 3µL DA responses across different session bins (mean ± s.d., Linear regression, R^2^ = 0.313, β = -0.566, *P* = 2.4e-5, *n* = 50 measures). G) Trial-average of the VS DA signal on 3µL patch in which mice only received the trial-initial reward but stayed for at least 3 seconds, experiencing repeated reward omissions. The DA signal “dips” below the baseline, to a lower level on rich patches, consistent with a larger rich value expectation (ΔDA, σ, mean ± SEM, *n* = 90 poor, *n* = 90 rich, see also **Figure S3F**). H) Comparison of the difference in 3µL omission responses across poor vs. rich patches (mean ± s.d., average between 1-3s, SR test, *P* = 0.0078, *n* = 10 mice). I) The difference between poor vs. rich patch omission responses across different time bins (mean ± s.d., Linear regression, R^2^ = 0.158, β = -0.346, *P* = 0.0147, *n* = 37 measures) J) Average VS DA responses to 3µL water rewards (pooled across poor and rich patches) received at different times on the patch (t = 1-5s). Responses to rewards increase and saturate with time on on patch, consistent with a diminishing effect of decaying value expectation **V_t_** (faded blue, see also **Figures S4G-H**). K) Correlation across mice between the difference in 3µL water responses across poor vs. rich patches (**E**), reflecting differential modulation by **V_t_**, and the rich environment PRT bias on 3µL patches, indicating a potential link between value expectation and PRT behavior (patch leaving decisions, Linear regression, R^2^ = 0.774, β = 5.44, *P* = 7.95e-4, *n* = 10 mice).

To test whether VS DA reflects a learned patch value function through a TD-RPE signal, we monitored VS DA across the full timeline of the foraging task using photometric recording of the DA fluorescence sensor GRAB_DA3m_^60^ (**Figure 4B**). We examined VS DA around specific events in the foraging task: **1)** transitions between blocks (entry into the poor or rich environment); **2)** rewards on rich vs. poor patches; **3)** omissions of expected rewards; and **4)** the amplitude of reward responses over time as value expectation diminishes (**Figure 4C**).

First, environment (i.e. block) transitions (**Figure S3A**) elicited positive VS DA responses, with a significantly larger response on rich environment entry (**Figure S3B-C**), as predicted from the TD-RPE model for transitions from low- to high-value states. The difference between poor vs. rich entry responses appeared over days (**Figure S3D**), indicating that mice learn different expected values for poor vs. rich environments which are not present in the task-naïve state.

If mice learn value functions that influence VS DA, we should also observe differences in responses to matched 3µL rewards across the two environments. Indeed, while DA exhibited positive responses for all rewards (**Figure S3E**), responses to 3µL rewards were larger on poor vs. rich patches (**Figure 4D-E**), consistent with negative modulation by reward expectation. Again, this difference was not present initially but emerged across days (**Figure 4F**), indicative of a learning process driven by experience of reward outcomes.

Reward expectations should also modulate VS DA responses when anticipated rewards are not received (omissions). Indeed, we also observed differential responses on trials in which probabilistic rewards beyond the first were not delivered (**Figure 4G**). DA decreased below the pre-trial baseline during omissions for all patches (**Figure S3F**) but decreased significantly more on 3µL omissions for rich vs. poor patches (**Figure 4H**), again emerging over days with a similar course as the reward response differential (**Figure 4I**). This indicates that VS DA is negatively modulated by reward expectations associated with poor vs. rich environment and patches, consistent with a TD-RPE signal influenced by a learned value function.

Finally, because patches in our task yield decaying probabilistic rewards (**Figure 1D, 3B**), the TD-RPE model makes the additional prediction that the amplitude of VS DA reward responses should increase with time on the patch, reflecting the time-varying (diminishing) modulation by value.^37,38,45,61,62^ To test for this signature, we examined DA responses on trials in which mice received only one probabilistic reward (in addition to the trial-initial reward) and quantified the response amplitude as a function of time (**Figure 4J**). DA reward responses scaled positively with time, mirroring the changing influence of expected value. The decay time constant ***τ*** of this inferred expectation signal (see **Methods**) was similar across environments and consistently followed an exponential shape across sessions (**Figure S3G-H**).

Taken together, these results show that the responses of VS DA during naturalistic patch foraging are consistent with a TD-RPE signal that reflects a learned value function. The fact that the differential poor vs. rich DA responses emerged over days, mirroring the evolution of the PRT bias (**Figure 2A, 2H, 3F**), suggests that this value function may directly influence foraging decisions. Indeed, across mice, the extent to which DA reflected the value differential was predictive of the PRT bias itself (**Figure 4K**), suggesting a potential causal relationship. However, as these data are only correlative, they do not indicate a causal role for DA as a learning mechanism or in shaping decisions. Thus, we next sought to test the hypothesis that DA drives construction of the value expectations that shape foraging decisions, leading to a PRT bias on rich patches.

### Dopamine reward signals are sufficient to produce a learned decision bias

To test the causal role of DA in this process,^29–35,63^ we used calibrated optical stimulation of DA neurons in the ventral tegmental area (VTA) to selectively boost phasic signals, creating artificially inflated reward responses (**Figure 5A**). In this experiment, mice navigated two environments with identical patch rewards and distributions (80% 1.5µL patches; 20% 3µL patches). However, in the fictive reward “FR” environment, we stimulated VTA DA neurons to boost the DA response during 1.5µL rewards to approximate a 6µL response (**Figure 5B-C**). Thus, while both environments carried identical objective reward outcomes, the manipulation was designed to produce fictive TD-RPE teaching signals. We hypothesized that this would shape construction of different value functions, biasing leaving decisions to drive a PRT bias in the FR environment (**Figure 5B**).

**Figure 5.**
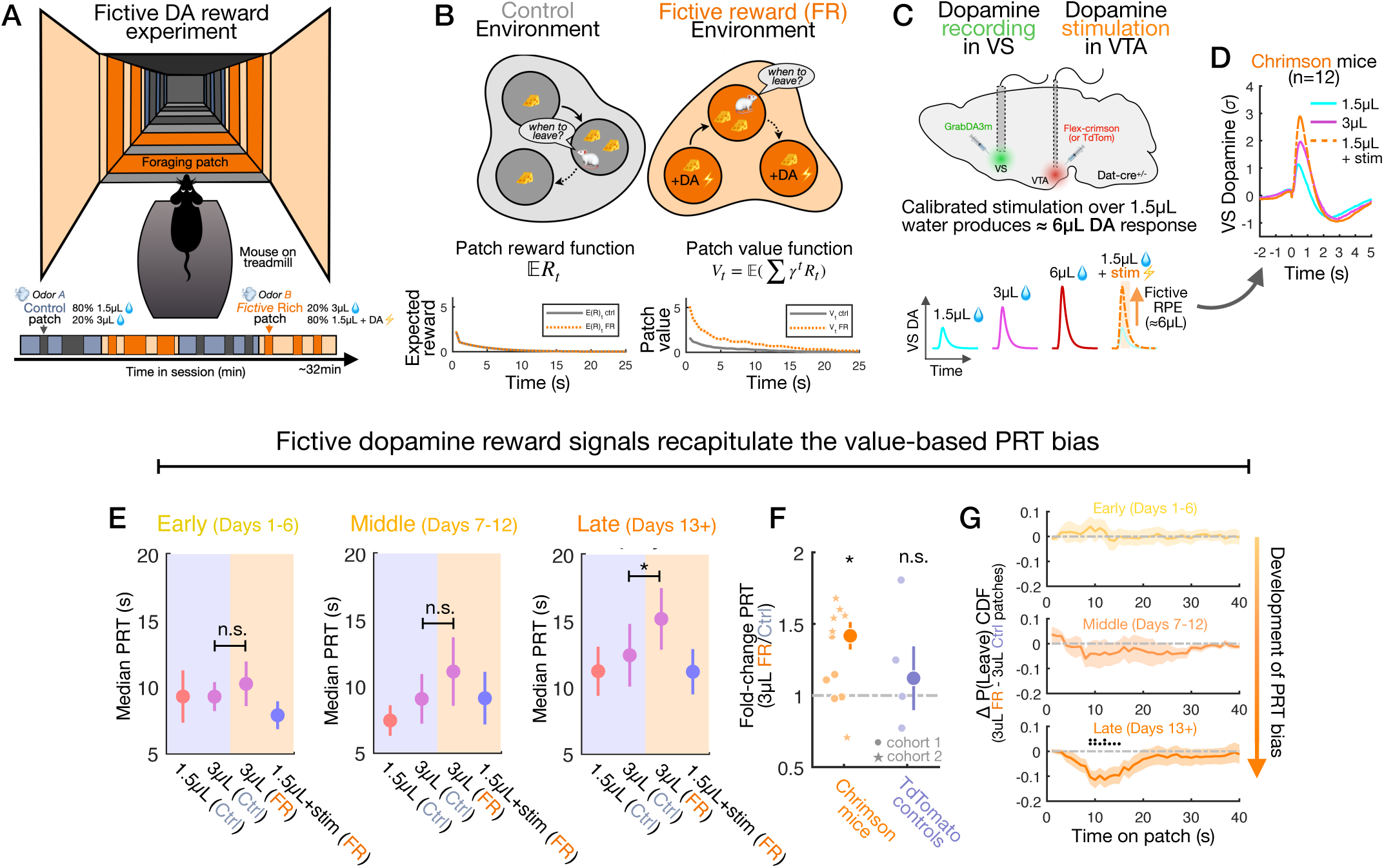
Phasic dopamine reward signals are causally sufficient for the value-based decision bias. **A)** Fictive reward (FR) experiment to test causal sufficiency of phasic dopamine for value learning and effect on each leaving decisions. Mice forage in two virtual environments with the same distribution of patches yielding 1.5µL (80%) or 3µL (20%) water rewards, but receive stimulation of dopamine (DA) neurons during 3µL rewards in the FR (orange) environment, boosting the reward signal to drive learning of fictive “rich” patch value. B) The expected reward functions E**R_t_** for FR vs. control patches are identical (left), but fictive reward signals are hypothesized to drive learning of different value functions **V_t_** (right) as with real 6µL rewards (Figure 3B). C) Experimental strategy to record ventral striatal dopamine (VS DA) using the GrabDA3m sensor while stimulating phasic physiological DA release using a cre-dependent excitatory opsin chrimson (or TdTomato) in dat-cre^+/-^ mice. Stimulation parameters (pulse duration and frequency) were calibrated per mouse before the foraging experiments to approximate a VS DA response of equivalent amplitude to a 6µL water reward (see **Figure S4, Methods**), creating a boosted reward signal. D) VS DA responses to 1.5µL (*n* = 1948 trials), 3µL (*n* = 862 trials), and 1.5µL + stim (∼6µL level, *n* = 2038 trials) in chrimson mice (*n* = 12). E) Patch residence times for 1.5uL, 3uL (control environment), 3uL (FR environment), and 1.5uL + stim patches across early (days 1-6), middle (days 7-12), and late (days 13+) sessions. Longer PRTs on FR 3µL patches (right) indicate a PRT bias driven by fictive DA reward signals on patches in the FR environment (median ± SEM, two sample t-tests, *P* = 0.39 [early], *P* = 0.26 [middle], *P* = 0.04 [late], *n* = 11 mice). F) The FR environment PRT bias on late days, expressed as the ratio of PRTs for FR vs. control 3µL patches, for stimulated (chrimson) and control (TdTomato) mice (median ± SEM, one sample t-tests, *P* = 1.27e-2, *n* = 11 mice [chrimson], *P* = 0.424, *n* = 4 mice [tdTomato]). G) Development of the PRT bias shown as the difference in P(Leave) CDFs for 3µL patches in the FR vs. control environment across early, middle, and late days.

We calibrated the stimulation by recording VS DA responses to rewards using photometry and tuning the parameters to reproduce the amplitude of natural 6µL responses (**Figure 5C**, **Methods**).^34^ Stimulation during 1.5µL water rewards in DAT-Cre^+/-^ mice expressing the excitatory opsin Chrimson^64^ approximated a 6µL reward signal (**Figure 5D, S4A**), while the same stimulation in tdTomato control mice produced minimal change in the 1.5µL signal (**Figure S4B-E**).

Mice were trained as before, and we monitored behavior and VS DA to test whether fictive reward signals could shape foraging decisions, analyzing PRTs across patch types and days. During early and middle sessions, we did not observe a significant difference in PRTs on 3µL FR vs. control patches (**Figure 5E**). Remarkably however, on late days, we observed significantly longer PRTs on 3µL FR patches which were never themselves stimulated (**Figure 5E**), recapitulating the positive PRT bias (**Figure 2A**). This difference was not present in control mice, ruling out the potential confound of optical stimulation alone (**Fig 5F**). We visualized the progressive development of the PRT bias by plotting the difference in leaving probability cumulative density functions (CDF, **Figure 5G**) across FR vs. control environments – while 3µL PRTs were indistinguishable early, mice exhibited a progressively stronger difference in PRTs over time.

This shows that phasic DA reward signals can drive value learning for self-paced foraging and shape foraging decisions. In addition to value learning, DA circuitry has also been variously implicated in the control of movement, learning dynamics, motivational states, and cognitive processes.^35,39–44,46^ While these diverse functions are not mutually exclusive, we sought to disambiguate the result of the FR experiment from other possible effects which could confound our interpretation. We analyzed movement of mice during stimulated vs. unstimulated rewards by measuring changes in velocity and acceleration of the running wheel and observed minimal effects (**Figure S5**), suggesting that while phasic DA signals can drive value learning, they do not strongly contribute to movement for self-paced foraging.^46,48,65^ In a subset of these mice, we also performed an additional experiment in which water rewards were completely replaced with different levels of DA stimulation (**Fig. S6A-D**). By itself, DA stimulation did not trigger movements or affect PRTs (**Figure S6E**), suggesting that it also does not immediately alter decision computations. Rather, this indicates that phasic DA reward signals in this context contribute specifically to value learning, shaping foraging decisions not through acute modulation of neural dynamics,^63^ but through their instructive role in forming value expectations that bias the decision process.

## Discussion

Foraging involves cognitive computations that balance reward gains (e.g. resources) against costs, providing an ecological window on value-based decision making – yet the neural basis of foraging decisions, and how they flexibly adapt in changing environments, is not well understood.^1–5^ Here, we studied mice making patch leaving decisions across environments with different global (poor vs. rich) and dynamic local reward statistics. Optimal theory predicts shorter time investments at rich-environment patches – but surprisingly, we observed the opposite: mice invested more time at rich patches with high expected yield, independent of actual reward outcomes, and this could be explained as an effect of value learning. Computational modeling of leaving decisions revealed a time-varying bias which mirrored the shape of a value function learned from reward outcomes. Consistent with a reinforcement learning (RL) mechanism,^21,22^ dopamine (DA) in the ventral striatum (VS) signaled a temporal-difference reward prediction error (TD-RPE),^23–27,45^ which could be used as a teaching signal to construct value functions, and calibrated manipulation of DA to create fictive TD-RPE signals recapitulated the value-related decision bias. These findings shed new light on the algorithms driving foraging behavior, expanding classical models by revealing a critical influence of value expectations on patch leaving decisions. More broadly, they establish direct evidence for DA-mediated value learning in ecological context, linking the canonical DA reinforcement learning framework to natural behavior and decision making.

### Value expectations shape patch leaving decisions across poor and rich environments

The fundamental computation underlying patch leaving decisions is a comparison of value across different scales: immediate reward gains are weighed against the potential for future long-term gains as an opportunity cost.^1,2^ The marginal value theorem (MVT, a classic optimality model) predicts that foragers should leave patches earlier in richer environments, because higher global reward rates raise this opportunity cost. We studied mice foraging across poor and rich environments, but we found that empirical behavior was not consistent with this. Instead, mice exhibited a systematic bias to invest more time (higher PRTs) at rich environment patches, specifically on late sessions once they had gained experience of reward statistics (**Figure 2A-C**).

Given this deviation from the MVT decision policy (**Figure S1**), which theoretically maximizes long-term reward gains, we wondered what could account for the rich PRT bias. Using a probabilistic decision model to measure the influence of reward variables on leaving decisions, we identified independent effects of reward outcomes and expectations driven by environment cues (**Figure 3F**). Within an environment, leaving decisions were oppositely modulated by immediate rewards and local reward history — that is, mice stayed longer when immediate outcomes were better, and left earlier when the recent reward rate was higher, indicating a strategy consistent with the MVT (**Figure 2I**). Yet across environments, learned patch values additionally biased mice toward staying at patches with high expected yields, limiting potential future gains by over-harvesting (consistent with similar observations in other foraging paradigms).^54,55^

Stepping away from the idealized MVT model, thinking about the natural context in which foraging takes place may put such apparent inefficiencies in a different light. Although this behavior results in fewer gains in the long run, it may also produce a more immediate benefit in the ecological setting: reduced sensitivity to missing expected rewards (i.e. disappointments) may allow animals to persist when they are locally “unlucky,” potentially harvesting more in the short term by waiting longer at a patch that their experience informs should be productive. This kind of systematic bias toward short term gains is also consistent with ideas about temporal discounting (animals tend to value future rewards less than immediate ones) and environment uncertainty. Theoretically, the MVT maximizes long term gains – but in the real world, animals do not have perfect information about reward distributions, rewards may be highly variable (imagine a mouse opening a nut to find it is rotten), and foraging cannot continue over infinite time. Animals may deal with the uncertainty and dynamism of the natural world by prioritizing the present, which is more certain and may be well informed by experience, such as with learned value estimates.

More broadly, the factors and algorithm governing how animals adjust cognitive strategies to account for, or to hedge against, the realities of real-world uncertainty (and update them with new informative experience) is an important general area for future research. It will also be valuable to test these models in other ecological task settings and species, including in humans, to gain insight into general principles and species-specific foraging strategies.

### Ventral striatal dopamine signals a TD-RPE during naturalistic foraging behavior

To elucidate the neural basis of value learning for foraging, we turned to an influential framework of biological RL (**Figure 4A**), which posits that animals construct value expectations from reward experience,^21,22^ with DA providing the critical teaching signal in the form of a TD-RPE.^24,27,34,45^ While the role of DA in reward learning is well established, this TD-RPE model has mostly been tested in simple association tasks, and it has been unclear how well it generalizes outside this domain. We recorded VS DA during self-paced foraging and observed several signatures of TD-RPE, including modulation of reward responses by poor vs. rich context, transients during state transitions, and scaling response amplitudes with diminishing value expectation. Consistent with other recent studies,^37,38,66^ this suggests that the TD-RPE model of DA is not explanatorily limited to conditioning or reinforcement paradigms but generalizes readily to more complex, naturalistic settings. VS DA also reflected the emergence of a learned value function through its influence on the TD-RPE signal (**Figure 4**), connecting DA-mediated value learning to foraging decisions.

Importantly however, we inferred this value function through its coordinated and correlated influence on decision behavior and VS DA (**Figure 4K**) – but our experiments did not address how value expectation is represented at the neural level, nor mechanistically how it alters foraging decision computations. Another recent study proposed that VS-projecting VTA DA neurons form a “minimal circuit” with recipient (and partially reciprocal) D1-receptor-expressing medium spiny neurons (D1-MSNs) to drive TD-RL in the context of classical conditioning.^34^ Although mechanistic details of this circuit remain to be clarified, the model raises two specific hypotheses relevant to the present study: that D1-MSNs encode value functions for patch foraging; and that the PRT bias produced by phasic DA stimulation is mediated by its effect on D1-MSN plasticity. Value representations in other areas, such as in prefrontal cortex, may also play an important role in flexibly adapting foraging decisions. In the future, these questions can be addressed using projection- and cell-type-specific recording modalities to track neural activity in specific circuit elements (e.g. D1-MSNs in VS) in concert with DA signals over the course of learning.

Our study also raises questions around the specific mechanisms underlying the influence of value expectations on decision computations. Recent studies, including our own,^11–13^ have shown that mice compute patch leaving decisions using a neural integrator decision variable (DV, akin to bounded accumulator dynamics for perceptual decisions),^6,14^ and these can be flexibly reconfigured across different reward contexts. The effect of value expectations on patch leaving decisions may be instantiated as a change in the speed (i.e. drift rate) of cortico-striatal ramping dynamics, effectively slowing the DV and increasing time to reach the leaving decision. Mechanistically, this could involve striatal circuits that both participate in integrator decision dynamics and receive input from dopaminergic TD-RPE signals (e.g. D1-MSNs).^67^ Other (non-exclusive) possibilities include cortical input to the DV (e.g. from prefrontal or retrosplenial cortex);^20,68^ thalamic input, which can modulate ramping dynamics for motor timing;^69^ and neuromodulators such as serotonin, which has been linked to patience for uncertain rewards.^18^

### Dopaminergic reward signals shape decision computations through value learning

Finally, despite the general success of the TD-RL framework to account for diverse properties of DA signals,^45^ the precise functionality of DA in the context of natural behavior remains intensely debated. Beyond reward learning and reinforcement, various models emphasize roles in the control of movement, learning dynamics, motivation, cognition, and threat assessment.^39,40,46,47,70–72^ There is also growing recognition that the activity of DA neurons is substantially more heterogenous than previously thought,^73,74^ suggesting that diverse functions may map onto distinct neural substrates. At the same time, specific hypotheses of DA function have largely been argued from correlative data, and rarely tested with precise, physiological manipulations in a naturalistic setting.

We found that artificially inflating phasic DA reward responses to create “fictive” TD-RPE signals was sufficient to recapitulate the value-based bias in foraging decisions (**Figure 5**), demonstrating a direct functional role in value learning. In additional experiments using similar manipulations (**Figure S5-6**), we found no evidence of immediate effects on movement or foraging decisions, suggesting that DA-mediated value learning can occur independently of motor or cognitive effects.^46,48,63,65^ These findings extend previous reports that DA stimulation can drive associative learning and behavioral reinforcement,^30–34^ demonstrating direct relevance of value learning in the context of foraging behavior. Importantly, our experiments targeted ventral tegmental area DA neurons non-specifically, and thus cannot disambiguate potentially distinct functions of different circuits (e.g. projections to ventral vs. dorsal striatum or frontal cortex).^73^ It will be critical in future studies to test physiological manipulation of projection-defined DA subpopulations, non-phasic stimulation, such as sustained “tonic” activation, and different magnitudes or schedules of rewards, which may play a role in shaping learning dynamics.^71,72^

Taken together, our results shed new light on the neurocomputational basis of foraging, revealing a novel role for value expectations in dynamic environments, establishing generalized function of biological TD-RL in ecological context, and demonstrating that canonical dopamine reward circuitry shapes foraging decisions by constructing value functions. The unification of this framework with foraging opens a powerful new avenue to elucidate principles and models of how animals learn and dynamically adjust cognitive strategies for real-world ecological problems.

## METHODS

### EXPERIMENTAL MODEL AND SUBJECT DETAILS

#### Mice

A total of 37 adult mice (20 male, 17 female, age 2-6 months) were used in the experiments: 20 C57/BL6j mice (9 male, 11 female) were used for the natural reward foraging task (**Figures 1-3**, **Figures S1-2**), and 10 (6 male, 4 female) of these were used for dopamine photometry (GRAB_DA3m_) recordings (**Figure 4**, **Figure S3**); 17 DAT-Cre^+/-^ mice (heterozygous for the gene expressing Cre recombinase under the control of the dopamine transporter DAT gene, 11 male, 6 female) were used for the optogenetics experiments (**Figure 5**, **Figures S4-5**, 12 experimental animals expressing Chrimson in dopamine neurons, and 5 control animals expressing tdTomato); of these Chrimson mice, those from the second cohort (5 males, 1 female) were additionally used for a second experiment in which we replaced water rewards with dopamine neuron stimulation (**Figure S6**). All mice were housed on a 12h dark/12h light cycle (dark from 07:00 to 19:00) and performed behavior tasks in the dark period. All procedures were performed in accordance with the National Institutes of Health Guide for the Care and Use of Laboratory Animals and approved by the Harvard Animal Care and Use Committee.

### METHOD DETAILS

#### Surgical procedures

For virtual-reality (VR) experiments, a custom titanium head-plate was attached to the skull for head-fixation. Mice were anaesthetized using isoflurane (4% induction, 1-2% maintenance) and a head-plate was attached to the skull with Metabond (Parkell). All surgeries were performed under aseptic conditions, and analgesia (Ketaprofin for post-surgery treatment, 5mg/kg buprenorphine for pre-operative treatment, 0.1mg/kg) was administered for 3 days following each surgery. The skin above the skull surface was removed and the skull dried with air, and we used a dental drill to create small craniotomies above virus injection sites. To express the third generation GRAB_DA3m_^60^ dopamine sensor in the ventral striatum (VS) for photometry, we unilaterally injected 300nL of AAV9-hSyn-GRAB-DA3m(DA3.3) (WZ Biosciences) into the VS (coordinates: +1.45mm [AP], 1.4mm [ML], -4.25mm [DV] from the skull surface), and then implanted a 400µm optic fiber (Doric lenses) 0.15mm above the injection site. To express the excitatory opsin Chrimson^64^ (or tdTomato) in the ventral tegmental area (VTA), we injected 250nL of flex-Chrimson (AAV8-hSyn-FLEX-ChrimsonR-tdT, UNC) or tdTomato (AAv5-CAG-tdTomato, UNC) unilaterally into the VTA (coordinates: -3.3mm [AP], 0.4mm [ML], -4.4mm [DV] from the skull surface) on the same side as the VS photometry fiber and implanted a 200µm 0.15mm above the injection site.

#### Foraging behavior experiments

For all experiments, mice were water restricted to maintain body weight > 85% of their baseline weight before training to ensure task motivation and maintain health. Additional water supplementation was provided following training sessions with a total of task and post-task water ranging from 0-3mL per day, determined individually per mouse. Mice that showed task motivation at higher weights were given greater amounts of water supplementation accordingly to minimize stress from water restriction. Body weights ranged from 85-105% of baseline.

#### Virtual reality (VR) patch foraging task

##### Setup for virtual reality foraging task

Virtual reality setups were identical to those used in our previous studies.^11,12,38^ Three monitors (width 53cm, height 30cm) were placed in front and on either side of the animal. Virtual reality scenes were generated using ViRMEn^75^ software in MATLAB on a workstation computer (DELL Precision 5810). Mice were head-fixed on a cylindrical Styrofoam treadmill (diameter 20.3cm, width 10.4cm) and the rotational velocity of the treadmill was recorded using a rotary encoder. The output pulses of the encoder were converted into velocity signals using custom Arduino code running on a microprocessor (Teensy 3.2) and integrated within ViRMEn to compute position. Water was delivered to mice from a waterspout placed in front of their mouth, and licks were monitored using an infrared sensor (OPB819Z, TT electronics). Voltage signals from the rotary encoder were digitized and recorded on the virtual reality computer using a data acquisition system (PCIe-6323, National Instruments). Water timing and amount was controlled using a solenoid valve (LHDA 1221111H, The Lee Company) and a switch (2N7000, On Semiconductor), with TTL pulses generated by the virtual reality computer via the PCIe-6323 data acquisition card.

##### Habituation

Mice were habituated to the treadmill prior to head-fixation with delivery of occasional water droplets from the reward spout while free to explore on top of the treadmill, which was held in place to prevent rotation. Once mice showed consistent interest in consuming water droplets (usually 2 days with 15-minute exposures), head fixation began on the following session. Initial head-fixation sessions were run for durations of 15 minutes, with gradual increases in session duration across consecutive sessions reaching a maximum duration of 50 minutes. During these sessions, water rewards (1.5µL, 2µL, or 6µL) were delivered in 10 second intervals with additional rewards given during running bouts to reinforce running behavior. Once starting head-fixation, mice performed daily sessions with 1 day off per week (6 days on, 1 day off) for the remainder of the task. Once mice were consistently running for most of the session (usually 3-5 days), they proceeded on the next session to the main foraging task.

##### Poor- and rich-environment foraging task

Once mice demonstrated consistent running behavior during habituation (above), they began training on the linear virtual track. Mice navigated through the virtual corridor and could stop at visually indicated patch locations to receive water rewards (they never received water between patches). The VR gain was set so that the travel distance between patches was 50cm. On patch entry, mice received an odor puff indicating that they were on the patch, and if they stopped moving for 0.05s-0.5s (incremented over days), could receive an initial water reward (t = 0s). Following this, they could stay on the patch to receive additional rewards each second with probability **P(Reward)** that decayed over time (**Figure 1D**). Initial **P(Reward) P_i_** at t = 1s was drawn randomly (uniformly) from **P_i_** = [0.15, 0.25, 0.58, 0.95] and followed an exponential decay with ***τ*** = 0.22s (***P_t_*** = ***P_i_e***^,***τt***^), with P_t_ = 0 after 30s.

Mice foraged in the poor and rich environment each session (30-40 min) in alternating blocks of approximately 7.5 minutes. Transitions between blocks was only triggered between patches and included a 10 second blank (black) screen before initiating the next block. The poor and rich environments were indicated with unique visual patterns on the corridor walls (between the patches), unique visual landmarks for patch locations, and a unique patch odor (isoamyl acetate or ethyl butyrate). In the poor environment, 80% of the patches yielded small 1.5µL rewards, and 20% yielded medium 3µL rewards; in the rich environment, 80% of the patches yielded large 6µL rewards, and 20% yielded medium 3µL rewards (**Figure 1E**). Within a given patch trial, all reward sizes were equal. The immobile time required to initiate the patch trial (starting at 0.05s) was gradually increased by 0.05s each day to a maximum of 0.5s. Occasionally, this was reduced for animals that showed faltering engagement rates and then increased again to 0.5s as engagement recovered. Mice exhibited positive scaling of patch residence times (PRT) with both reward size and frequency (**Figure 1F**), indicating strong sensitivity to immediate reward outcomes.

##### Fictive reward dopamine stimulation experiments

We performed two additional experiments using optogenetic stimulation of dopamine neurons to produce fictive reward signals. We first tested the effects of using calibrated stimulation to artificially inflate responses to real 1.5µL water rewards (**Figures 5, S4-5**), producing responses equal in amplitude to 6µL rewards in the “fictive rich” (FR) environment. We trained two cohorts (*n* = 17) of DAT-Cre^+/-^ mice in this task. The virtual reality setup, visual stimuli, odor cues, and task structure (reward statistics, block lengths, etc.) were identical to the natural reward task described above, except that the FR environment “large” patches yielded 1.5µL instead of 6µL rewards, and optical stimulation of VTA dopamine neurons was applied during all FR 1.5µL rewards. The stimulation duration was 0.4s (during rewards), and parameters (pulse frequency, pulse length) were calibrated for each mouse based on free 6µL rewards given before the task during the habituation period (**Table S2**).

We performed a second experiment (**Figure S6**) in a subset of mice (*n* = 6 DAT-Cre^+/-^ animals expressing Chrimson in VTA dopamine neurons) to test the effects of dopamine stimulation alone (not coupled with real rewards). After the first experiment, these mice were given a break of approximately three weeks in which they had no task exposure and then were trained on the second task. In this task, patches always yielded one trial-initial real reward (6µL a t = 0s), followed by 0.4s stimulation with exponential decaying probability (***τ*** = 0.22) each second to create reward-like dopamine transients in place of water rewards. Stimulation was delivered at three frequencies to create three different amplitudes approximately scaled to the level of 1.5µL, 3µL, and 6µL reward responses (**Table S2**). The task structure (reward probabilities, patch distributions, session and block lengths) was otherwise the same as in the previous task, except a novel set of odor cues and visual patterns for the virtual environments and patches were used.

#### Histological analysis and validation of fibers and virus expression

Once experiments were completed, mice were transcardially perfused with phosphate buffered saline (PBS) followed by 4% paraformaldehyde in PBS. The brains were sliced in 100 µm coronal sections using a vibratome (Leica). Brain sections were mounted on glass slides and stained with 4’, 6-diamidino-2-phenylindole (DAPI, Vectashield) in mounting medium. Slides were then imaged with a fluorescence microscope (Zeiss, Axio Scan.Z1). Localized expression of the dopamine fluorescence sensor GrabDA3m (DA3m), and placement of optic fibers above the recording site in the ventral striatum (VS), were visualized and validated for each mouse (example, **Figure 4B**). Expression of the Cre-dependent excitatory opsin flex-Chrimson (or tdTomato) in the ventral tegmental area (VTA) of DAT-Cre^+/-^ mice, and placement of optic fibers above VTA, were confirmed histologically (**Figure S4A**).

#### Mouse and patch trial inclusion criteria

A total of 37 mice were trained in the natural reward or the fictive reward foraging tasks. In virtual-reality head-fixation, mice sometimes disengaged from the task by pausing for brief periods of time, and this occasionally occurred on a patch trial. To prevent contamination of decision behavior by this disengaged state, we did not include trials with patch residence time greater than 60 seconds for behavior analysis (594/23084, 2.57% of all trials dropped). For analysis of foraging decisions across environments (**Figure 2A-C**, **Figure 5E-G**), we required mice to perform at least 10 medium 3µL trials to measure and compare PRT behavior. All mice met this inclusion criterion for the natural reward experiment. For the fictive reward optogenetic experiments (**Figure 5**), one Chrimson mouse and one tdTomato control mouse became disengaged from the task and did not perform enough trials to meet this criterion.

#### Analysis of patch residence time (PRT) decision behavior

We analyzed decision behavior for patch foraging trials using the patch residence time (PRT, the time mice stay at a given patch trial) for different trial types. To evaluate whether mice made patch leaving decisions in a value-sensitive manner, we compared PRTs across patches with different reward sizes, and for each reward size, across hidden trial types that had different initial reward probabilities (P_i_ starting at t = 1s, after the initial reward). P_i_ was not cued and was drawn randomly on trial initiation (see task descriptions). Mice were sensitive to both reward size and frequency, staying longer on patch trials with better reward outcomes (**Figure 1F**, **Figure 2F** immediate reward regressors, **Figure 3F** weight on patch reward rate).

To quantity the effect of poor vs. rich environment (**Figure 1B**) on patch leaving decisions, we compared PRTs across matched, medium 3µL patches in the poor vs. rich environment. On late sessions (days 13+), PRTs on rich environment 3µL patches were significantly higher than on poor environment 3µL patches (**Figure 2A-B**). In part to rule out the potential confound of local reward rate (both in the session and on recent patch trials), we performed a multiple regression analysis including a set of regressors capturing reward information (**Figure 2F**). While immediate rewards and local reward history influenced patch leaving decisions, the effect of rich vs. poor environment identity was still significantly positive, indicating a rich environment bias independent of immediate or local reward experience.

We also quantified motivated behaviors in the foraging task to test whether mice behaviorally distinguish the poor vs. rich environment outside of patch leaving decisions (**Figure S2**). We quantified patch engagement as the fraction (%) of patches that mice stopped at for long enough (the immobile time, 0.05-0.5s) to engage the trial and receive the trial-initial reward. For individual mouse engagement measures, we quantified this for each session and took the average over sessions. We also quantified the speed with which mice moved between patches (the inter-trial-intervals). We identified pairs of successively engaged patch trials within an environment, measured the time from patch leave to the next patch entry, divided 50cm (the distance between patches) by this time, and averaged these within each mouse.

#### Multiple regression model of patch residence time against reward variables

To quantify the varied effects of reward variables on patch leaving decisions, we performed a multiple regression analysis, in which trial PRTs were modeled as a linear weighted sum of task variables (**Figure 2D**). For each mouse, we formed a regressor matrix **X** by pooling their patch trials across sessions and computing 9 reward-related and 4 temporal variables associated with each trial. These included: session day; the time in session; the time since the last block switch; the time since the last patch trial; the local reward rate history (reward rates on patches n-1, n-2, n-3, n-5, n-10) and session reward rate; immediate reward frequency and size on the current patch; and rich vs. poor environment identity, which captures the effect of cue-driven reward expectation. For each mouse’s model, we normalized the regressor matrix and outcome variable (trial PRTs) by z-scoring, so that the coefficients **β** were interpretable as directional change in PRT in units of *σ* (**Figure 2F**). To quantify effect sizes in seconds (**Figure 2G**), we re-scaled **β** by the standard deviation of the PRT distribution used as the outcome variable for that mouse’s model.

Models were validated by comparing the performance of the full model in capturing PRT variance (R^2^) against a model constructed with shuffled outcome variables, and by comparing the performance to predict held-out data (10-fold cross validation) against null models with shuffled data (**Figure 2E**). Effects of reward information on PRT were quantified as the average coefficient **β** across models for all mice. To quantify regressor time-courses over sessions, we concatenated trials across mice and sessions for each time bin (successive 3-day bins) and then averaged across coefficients **β** for different types of reward information (e.g. immediate rewards, local reward rate history, **Figure 2H**).

#### Time-varying probabilistic patch leaving decision model

To gain insight into how patch leaving decisions are modulated in real time, we constructed a decision model that predicts a time-varying patch leaving probability **P(Leave)_t_** (1 - **P(Stay)_t_**) based on immediate reward outcomes (on the patch), recent reward history, and a time-varying environment-specific bias (**Figure 3D**). The model takes a logit threshold (with temperature parameter ***T***) over a weighted linear combination of these factors: the current patch reward rate at time t on the patch (**∑R/t**), weighted by **ƛ**; the reward rate history from recent patches (with an exponential decaying weight ***τ*** = 0.063 on patches n-1, n-2, n-3, n-5, and n-10, with ***τ*** fit from the equivalent coefficients from the PRT regression analysis), weighted by ***ρ***; and a set of biases ***β_t_*** for poor and rich patches tiling each second on the patch. The model parameters ***T***, **ƛ**, ***ρ***, and ***β_t_*** for trials pooled across sessions for each mouse were fit using maximum likelihood estimation (*fmincon* in MATLAB), and model performance was quantified using the negative log likelihood of predicted vs. real patch leaving probabilities on holdout trials (10-fold cross validation). We compared the full model with environment-specific biases (Kernel) ***β_t_*** against reduced models with a single constant bias (Flat) or no bias (None, **Figure 3E**).

We analyzed the fits of the model parameters (for full models) across session bins to understand the relative influence of these factors on patch leaving decisions (**Figure 3F**). While the bias parameters ***β_t_*** were flat on early days, indicating a constant bias, an exponential decay structure emerged over time, which was higher amplitude for rich patches. We analyzed the shape of ***β_t_*** by fitting an exponential function and comparing the amplitudes of fit exponentials across time and poor vs. rich patch ***β_t_*** (**Figure 3H**).

#### Analysis of patch leaving decisions using optimal foraging theory

To gain intuition for how mice might make patch leaving decisions in poor vs. rich environments, we simulated patch leaving behavior under a decision policy based on the Marginal Value Theorem (MVT).^1,2^ A foundational result in optimal foraging theory, the MVT gives the optimal leave time for patch foraging (in the sense of long-term reward maximization) as the time when the marginal (in the limit instantaneous) reward rate equals the long-term future expectation (the global reward rate, **Figure S1A**). Intuitively, it is optimal to leave when the marginal gain from staying an additional small time (i.e. **δ**_reward_ in **δ**t) is no longer worth the time opportunity cost compared to the long-term reward rate expected from leaving the patch (Δ_reward_ in Δt). To implement an MVT-based policy, a forager must therefore use estimates of both the immediate on-patch reward rate (**δ**_reward_/**δ**t) and the long-term global reward rate (Δ_reward_/Δt).

To simulate MVT decision behavior for our task, we assumed that mice could estimate environment reward rates from their experience, which we computed as 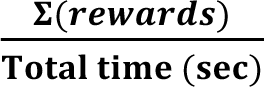 for all patch trials in a session (the global reward rate), or for each environment (the poor and rich environment reward rate). We took the average of these quantities across mice and sessions (**0.1127** rewards/second for the poor environment, **0.4645** rewards/second for the rich environment, and **0.2863** rewards/second for the global reward rate) to be an empirical measure of long-term reward rates mice could potentially use, and we simulated leaving decisions (described below) against these values to generate leaving time (PRT) distributions under the global reward rate policy or environment-specific reward rate policy.

For probabilistic rewards without reward expectations, mice must rely on ongoing estimates of the local reward rate **δ**_reward_/**δ**t on each patch trial to make leaving decisions. We assumed that mice could approximate **δ**_reward_/**δ**t using the local objective reward rate, which they experience on each trial. We computed this in a rolling 3-second window for 1000 trials of each patch type (1µL, 3µL poor, 3µL rich, 6µL), and for each trial, compared it against Δ_reward_/Δt to generate simulated patch leaving decisions. As expected, when compared against the same global reward rate, MVT-simulated PRTs were indistinguishable for 3µL poor vs. 3µL rich trials (**Figure S1B-C**). When using environment-specific reward rates, simulated PRTs were significantly shorter in the rich environment (**Figure S1D-E**). This demonstrates the logic and theory of the MVT: rich-resource environments represent a higher opportunity cost for staying at any given patch, so the MVT predicts shorter PRTs for reward-matched patches in rich vs. poor environments.

#### Temporal-difference reinforcement learning of patch value functions

To understand how a patch value function might emerge through biological reinforcement learning (RL), we simulated the development of a value function **V_t_** by applying the temporal-difference (TD) learning algorithm^21,22^ (**Figure 4A**) to empirical reward returns experienced by mice in our foraging task. The TD learning algorithm iteratively constructs an estimate of a value function **V_t_** which represents an expectation of current and future reward returns over a discounted time horizon: ***V_t_*** = E(**Σ*γ^t^R_t_***), where **γ** is the discount factor, between 0 and 1. As rewards **R_t_** are experienced, **V_t_** is updated by the TD reward prediction error (TD-RPE, or TD error ***δ_t_*** using the update rule ***V_t_*** ← ***V_t_*** + ***αδ_t_***, where ***α*** is the learning rate). The TD error ***δ_t_*** measures deviation between experienced and expected reward (***δ_t_*** = ***R_t_***_+**1**_ + ***γV_t_***_+**1**_ − ***V_t_***), and so through successive updates, converges to the value function **V_t_** which minimizes reward surprise.

We applied the TD learning algorithm to empirical reward returns on foraging patches (**Figure 3B**, ***α*** = 0.03, **γ** = 0.8) and plotted the resulting estimates of the value function **V_t_** after 5 trials (early), 10 trials (middle), and 100 trials (late). We then measured how **V_t_** for poor and rich patches changed over learning by fitting exponentials to simulated **V_t_** (**Figure 3C**) to guide specific predictions for how the PRT decision bias may change over time under the hypothesis that it is driven by a learned value function.

#### Photometric recording of dopamine release in the ventral striatum

Fluorescence signals were recorded using a custom-made fiber photometry system as described in our previous studies.^37,38^ The blue light (473 nm) from a diode-pumped solid-state laser (DPSSL; 80–500 μW; Opto Engine LLC, UT, USA) was attenuated through a neutral density filter (4.0 optical density, Thorlabs, NJ, USA) and coupled into an optical fiber patchcord (400 μm, Doric Lenses) using a 0.65 NA microscope objective (Olympus). The patchcord connected to the implanted fiber was used to deliver excitation light to the brain and to collect the fluorescence emission signals from the brain. The fluorescent signal from the brain was spectrally separated from the excitation light using a dichroic mirror (T556lpxr, Chroma), passed through a bandpass filter (ET500/50, Chroma), focused onto a photodetector (FDS100, Thorlabs) and amplified using a current preamplifier (SR570, Stanford Research Systems). The voltage signal from the preamplifier was digitized through a data acquisition board (PCI-e6321, National Instruments) at 1kHz and stored as a task variable in MATLAB running the virtual-reality software (ViRMEn).

#### Analysis of dopamine signals around events in the foraging task

The raw dopamine photometry signal for each session was pre-processed by first smoothing the signal using a 50ms left moving average and then normalized by z-scoring the full session trace. Activity was then extracted trial-by-trial for several task events and aligned to their onset: *environment entry* (when a block transition occurred and the new environment appeared on the screen); *patch entry* (when the mouse entered the patch location); and *patch trial onset* (when the trial-initial reward was received).

To quantify responses to reward events (receipts and omissions, **Figures 4D, G, J**), we further normalized single-trial signals by subtracting the immediate pre-trial baseline (−0.1:0s from the event time). To quantify responses to the block transitions (**Figure S3B**), we subtracted a longer baseline window (−5s:0s from the event time, during the blank screen in between virtual environments). Quantification of dopamine responses throughout are reported as the change from pre-trial baseline in units of standard deviation (i.e. ΔDA *σ*). For all plots of trial-averaged dopamine signals, the mean ± SEM (standard error of the mean) across trials is shown.

To measure the change in dopamine reward response amplitudes with time on the patch (**Figure 4J**), we averaged over trials in which mice received only one reward at a specific time (in addition to the trial-initial reward) to eliminate contamination from other reward responses. To measure the decay of an inferred value function which may drive this time-varying modulation, we reflected the peak responses around their midpoint (see faded blue trace in **Figure 4J**) and fit an exponential decay function. This was well fit by an exponential across early, middle, and late days (**Figure S3H**). The decay constant ***τ*** was nearly identical across poor (***τ*** = 0.266) and rich (***τ*** = 0.263) patches (**Figure S3G**), and close to the value of the decay constant used to generate exponentially decaying reward probabilities (***τ*** = 0.22, **Figure 1D**), indicating a close match to the hypothesized value function.

#### Calibration of optogenetic stimulation to produce fictive phasic reward signals

For the fictive reward foraging experiments, we calibrated optical stimulation of VTA dopamine neurons to closely reproduce either the amplitude and dynamics of a 6µL ventral striatum dopamine (VS DA) water reward response when stimulation was applied during a 1.5µL reward (experiment 1), or to produce reward-like phasic responses at three different amplitudes in the absence of any reward (experiment 2). Calibration was performed for each mouse prior to the task. During the habituation phase, when mice were given free water rewards, we gave 6µL rewards and 1.5µL rewards coupled with optical stimulation of VTA (power = 3mW (∼24mW/mm^2^) at fiber tip, duration = 0.4s)^34^ with different parameter combinations (pulse length = [8ms – 12ms], pulse frequency = [5Hz – 30Hz]), and over 1-3 sessions, we identified parameters that closely reproduced the 6µL response (see example mouse 948, **Figure S4C**, and group-level average responses, **Figure 5D**). Once determined, the parameters for each mouse were set for the entire experiment. For the second experiment, we used 10ms pulses at 6Hz (small), 14Hz (medium), and 25Hz (large) for all mice, as this consistently generated three amplitudes of reward-like dopamine transients in the same range as responses to 1.5µL, 3µL, and 6µL water rewards (**Figure S6**). Control tdTomato mice were stimulated using parameters that typically produced ∼6µL reward response amplitudes in Chrimson mice (see **Table S2**).

### QUANTIFICATION AND STATISTICAL ANALYSIS

All data were analyzed with custom code written in MATLAB (version 2019b, Mathworks). For statistical tests, normality of data and equal variance of groups were not assumed by default, and non-parametric tests were used when these criteria were not met. Tests for normality were used before applying any parametric tests (t-tests, ANOVA), and for all significance testing we used two-tailed tests at α = 0.05 or α = 0.01 (noted in figure legends). All 95% confidence intervals are reported for test statistics and effects, and when necessary, were estimated by re-sampling. For the PRT regression analysis and the probabilistic decision model, performance was quantified by measuring predictions on held-out trials (10-fold cross-validation, using R^2^ or -log likelihood). The sizes of mouse groups were not pre-specified and approximated those of previous studies. Comprehensive details for all statistical tests are provided in **Table S1**.

## ACKNOWLEDGEMENTS

We thank Edward Soucy and Yuwei Li of the Harvard Neurotechnology Core for engineering assistance; the Harvard Center for Biological Imaging for support with microscopy; and Mitsuko Watabe-Uchida and Uchida lab members for discussion and critical feedback. This work was supported by grants from the National Institutes of Health (NIH) (5U19NS113201 to N.U. and 5R01DA05975 to N.U.) and the Simons Collaboration on Global Brain (to N.U.). L.K. was supported by a Helen Hay Whitney Fellowship.

## AUTHOR CONTRIBUTIONS

L.K. designed the foraging tasks and built experimental setups. L.K. and H.M. performed behavior and recording experiments for the natural reward task and the fictive reward tasks. L.K. and H.M. analyzed behavior, dopamine physiology, and computational models. L.K. wrote the manuscript with feedback from N.U. No generative AI tools were used for any aspect of this work.

## DATA AND CODE AVAILABILITY

All code and data will be uploaded to a public repository upon publication.

---

**Figure S1.**
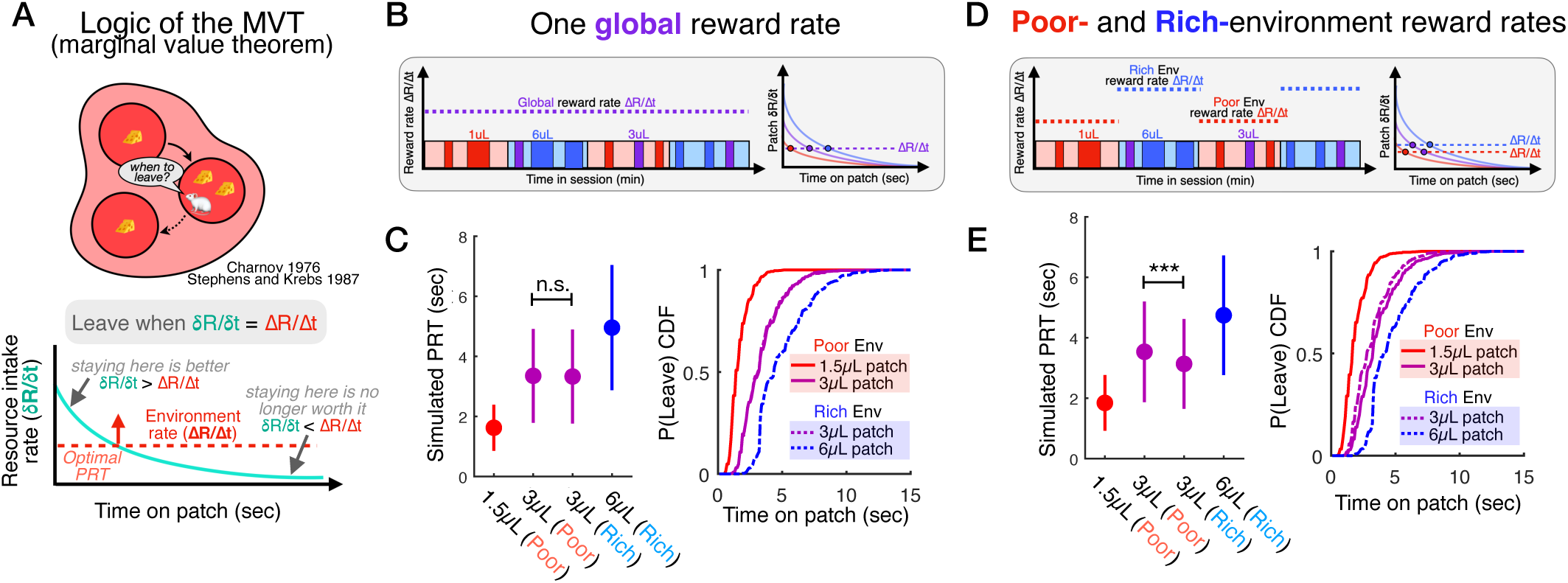
The marginal value theorem predicts a negative patch leaving bias in the rich environment. **A)** Cartoon illustration of the logic of the marginal value theorem (MVT, Charnov 1976), which states that the optimal leave time for a foraging patch occurs when its marginal reward rate falls to the level of the long-term environment reward rate. **B)** Simulation of patch leaving decisions across poor-vs. rich-resource environments using the same patch structure and reward statistics as in the mouse foraging task by applying an MVT decision policy (see Methods) and assuming that mice compare marginal patch reward rates to a single global reward rate across environments. **C)** PRTs for simulated MVT decision behavior under a single global threshold for small (1.5µL), medium (3µL), and large (1.5µL) reward patches, comparing PRTs across reward-matched poor vs. rich environment patches. The single global reward rate threshold leads to identical patch leave time distributions (mean ± s.d., *P* = 0.7476, two sample t-test, *n* = 1000 simulated trials for each patch type). **D)** Simulation of patch leaving decisions by applying an MVT decision policy (as in **B**), except assuming that mice compare marginal patch reward rates to poor- or rich-environment-specific global reward rates. **E)** PRTs for simulated MVT decision behavior under the environment-specific reward rate policy, comparing PRTs across reward-matched poor vs. rich environment patches. The high reward rate in the rich environment leads to earlier patch leave times in the rich environment (i.e. a negative PRT bias, mean ± s.d., *P* = 1.5e-8, two sample t-test, *n* = 1000 simulated trials for each patch type).

**Figure S2.**
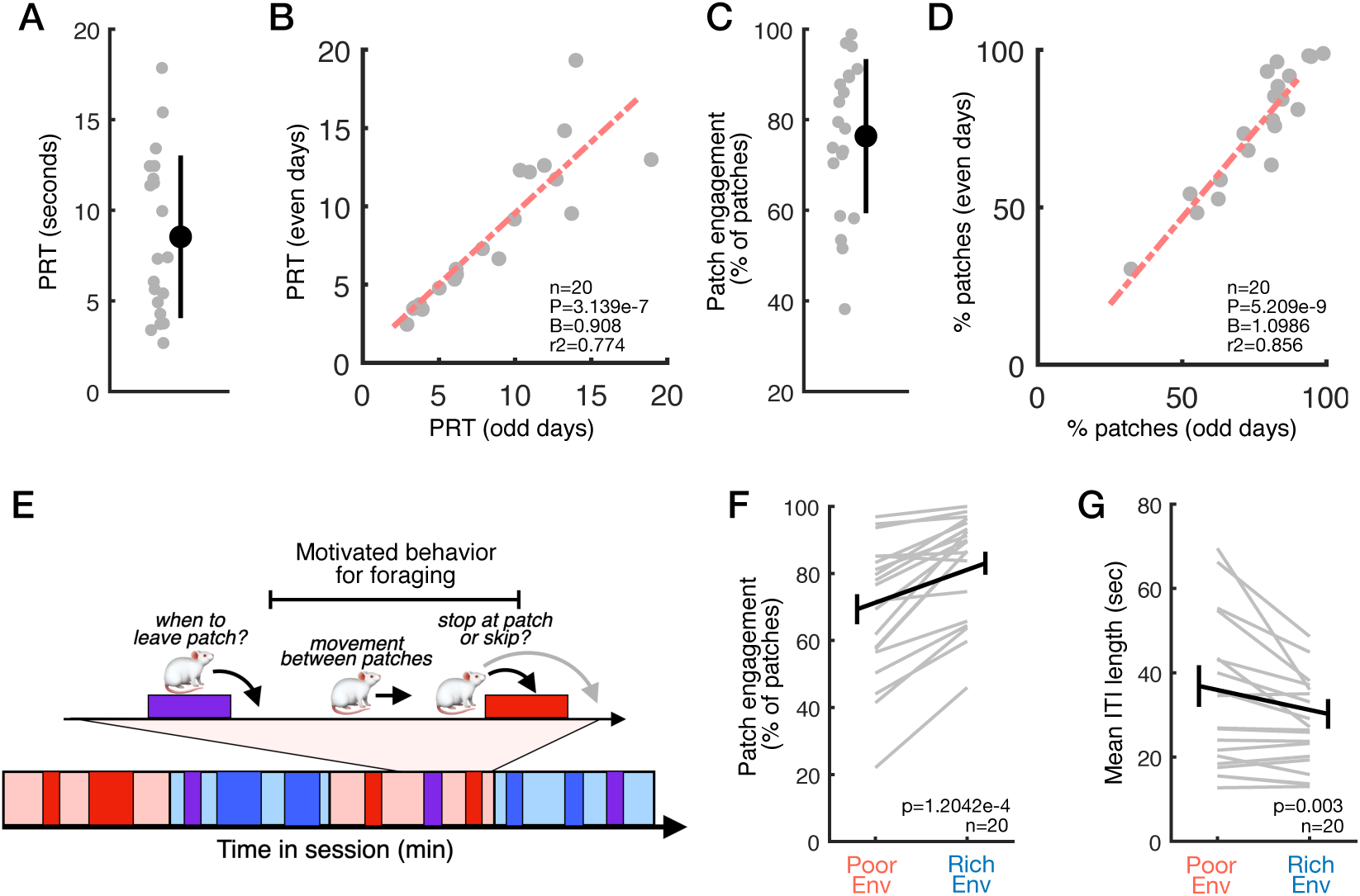
Additional detail and analysis of mouse behavior in the virtual patch foraging task. **A)** Median patch residence time (PRT) for each mouse (each dot) across all patches, showing group-level mouse average and variability (mean ± s.d., *n* = 20 mice). **B)** Stable individual variability in PRTs, shown and quantified as the linear correlation between median PRTs across session splits (odd and even days, R^2^ = 0.856, β = 0.908, *P* = 3.139e-7, *n* = 20 mice). **C)** Overall patch engagement (the fraction of patches visited) for each mouse (each dot) across all patches (mean ± s.d., *n* = 20 mice). **D)** Stable individual variability in patch engagement, shown and quantified as the linear correlation of % patches visited across session splits (odd and even days, R^2^ = 0.774, β = 1.098, *P* = 5.21e-9, *n* = 20 mice). **E)** Cartoon illustrating measures of motivated behavior in the foraging task. Patch engagement (the fraction of patches visited) and movement between patches are independent of individual patch leaving decisions and reflect mice’s motivation to forage for water rewards in each environment. **F)** Comparison of patch engagement across poor vs. rich environments. Mice are more likely to stop and harvest rewards from patches in the rich environment (*P* = 1.2e-4, SR test, *n* = 20 mice). **G)** Comparison of movement (inter-trial interval [ITI] times across poor vs. rich environments. Mice move more quickly between successively engaged patches in the rich environment (*P* = 3.2e-3, SR test, *n* = 20 mice).

**Figure S3.**
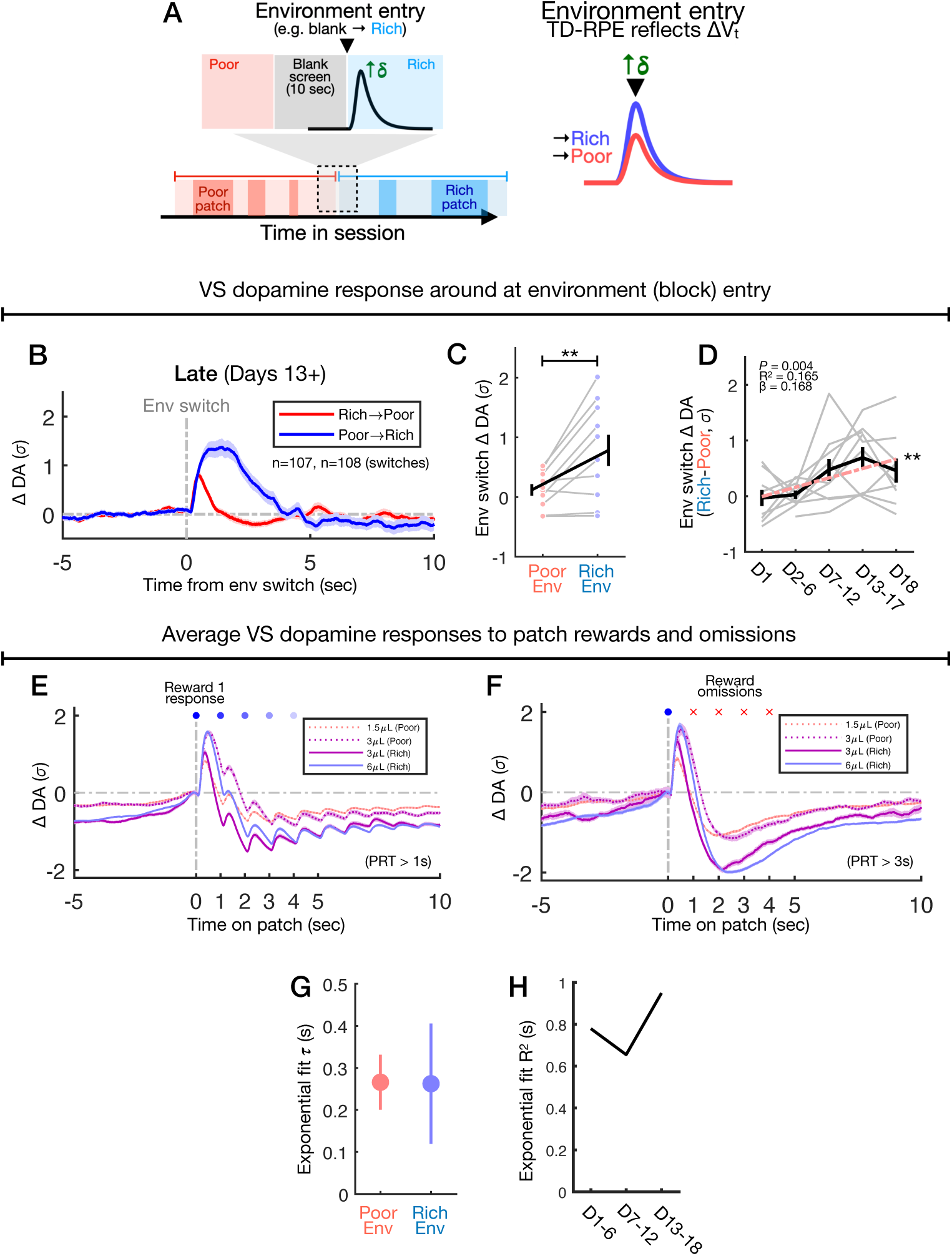
Additional analysis of ventral striatal dopamine responses to events during foraging. **A)** Schematics illustrating the environment entry event in the virtual foraging task. When a block ends (after approximately 7.5 minutes), the virtual-reality screen is blank for 10 seconds, and then the alternate environment is shown (the screen turns back on). If dopamine signals a temporal-difference reward prediction error (TD-RPE, δ**_t_**), then it should exhibit a positive transient on environment entry, reflecting a change from a lower value (the blank screen) to higher value (the virtual environment) state (i.e. **ΔV_t_** > 0, and δ**_t_** > 0). **B)** Response of ventral striatal dopamine (VS DA) around entry events into the poor (red) or rich (blue) environment on late sessions (days 13-18). The DA signal is expressed as change from the pre-trial baseline period in z-scores (ΔDA, σ, mean ± SEM, *n* = 107 poor entry events, *n* = 108 rich entry events) **C)** Comparison of VS DA poor vs. rich environment entry responses on late sessions across mice (mean ± s.d., SR test, *P* = 0.0098, *n* = 10 mice) **D)** Difference in entry responses (**C**) across time bins from early to late days showing development of poor vs. rich differential over time (each line a mouse, mean ± s.d., Linear regression, R^2^ = 0.165, β = 0.168, *P* = 0.0042, *n* = 48 measures). **E)** Average responses of VS DA to water rewards for all patch types (1.5µL, 3µL poor, 3µL rich, 6µL), aligned to the first reward at t = 0s and normalized to the local pre-trial baseline (ΔDA, α, mean ± SEM, *n* = 1404 1.5µL, *n* = 362 3µL poor, *n* = 405 3µL rich, *n* = 1646 6µL). **F)** Average responses of VS DA on trials in which mice did not receive rewards beyond the first (omissions, X) but stayed on the patch for at least 5 seconds. Responses for all patch types (1.5µL, 3µL poor, 3µL rich, 6µL) are aligned to the reward at t = 0s (ΔDA, σ, mean ± SEM, *n* = 375 1.5µL, *n* = 90 3µL poor, *n* = 90 3µL rich, *n* = 476 6µL). **G)** Measurement of time-varying VS DA water response amplitudes for patches in the rich and poor environment. The trial-averaged response amplitudes (Figure 4J) were reflected to fit an exponential decay time constant (τ) of the inferred diminishing effect of value **V_t_** on the VS DA response with time on the patch (τ, 95% CI). **H)** Performance measure (R^2^) for an exponential fit to the reflected reward response amplitudes over time (**G**) for different time bins (early days 1-6, middle days 7-12, late days 13+).

**Figure S4.**
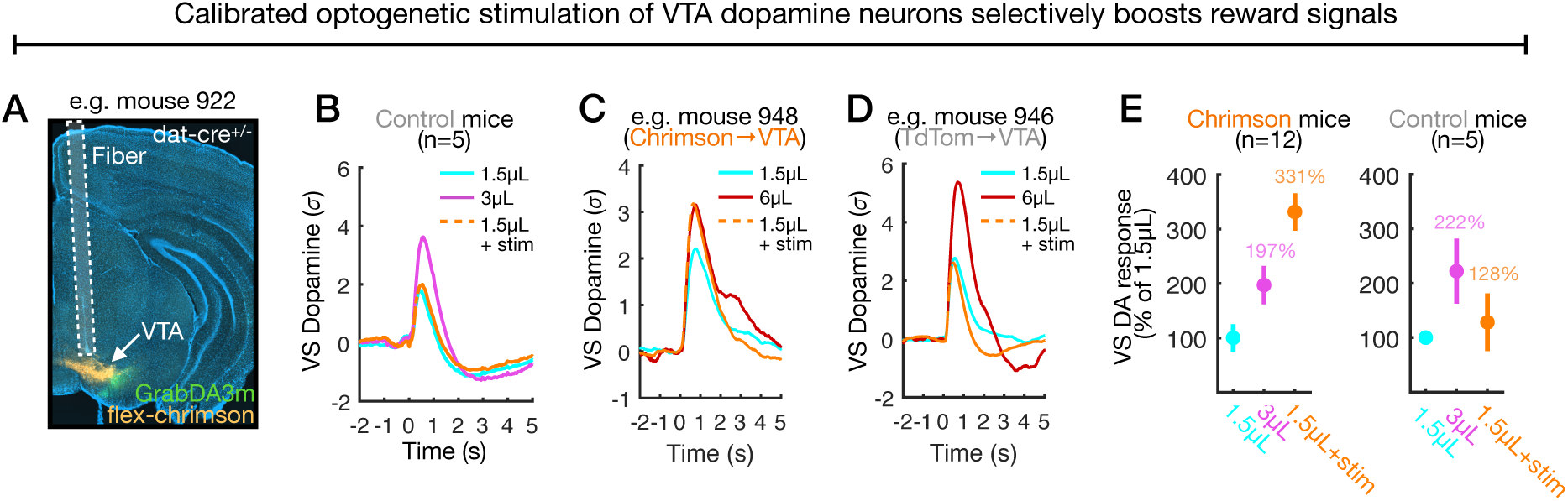
Supporting data and validation of calibrated optical stimulation of dopamine neurons. **A)** Histology image from example mouse 922 showing expression of the cre-dependent excitatory opsin (flex-chrimson) in midbrain (ventral tegmental area, VTA) dopamine neurons (orange) and placement of optic fiber. **B)** VS DA responses to 1.5µL (*n* = 446 trials), 3µL (*n* = 198 trials), and 1.5µL + stim (*n* = 417 trials) in tdTomato mice (*n* = 5). **C)** Ventral striatal dopamine (VS DA) responses to natural water rewards (1.5µL, 6µL) and 1.5µL rewards coupled with calibrated optogenetic stimulation of VTA DA neurons to produce “boosted” fictive responses at the level of a 6µL reward response (example dat-cre^+/-^ mouse 948 expresses chrimson in VTA DA neurons). **D)** VS DA responses to natural water rewards (1.5µL, 6µL) and 1.5µL rewards coupled with optogenetic stimulation in an example control mouse 946 expressing td-Tomato, showing no change in the signal on stimulated trials. **E)** The average amplitude of VS DA responses to natural water rewards (1.5µL, 3µL) and 1.5µL rewards coupled with optogenetic stimulation, normalized to the 1.5µL response amplitude, for chrimson mice (*n* = 12, left) and tdTomato mice (*n* = 5, right). The amplitude of the fictive reward response generated with calibrated VTA DA stimulation is approximately 231% higher than the 1.5µL response (mean ± s.d.).

**Figure S5.**
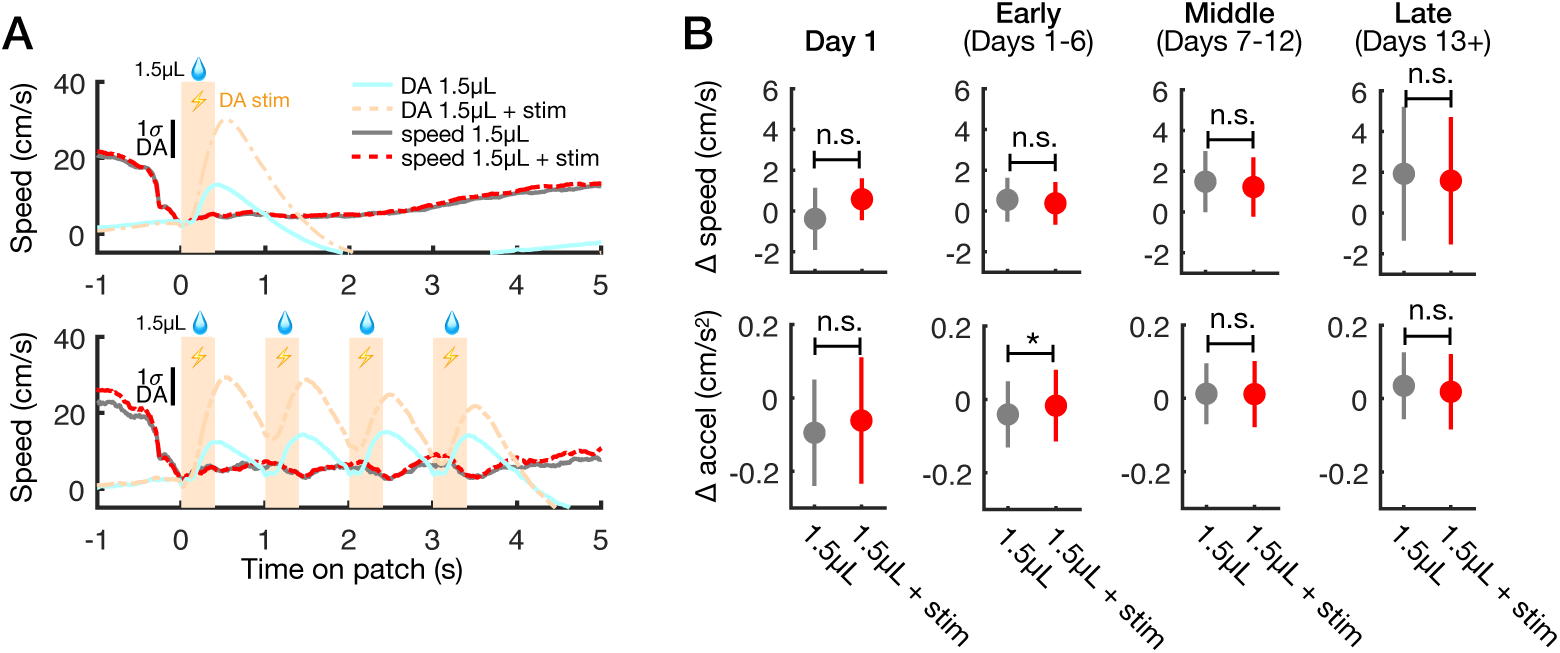
Immediate effects of phasic dopamine stimulation on mouse movement. **A)** Measurement of mice’s movement on the running wheel (speed in cm/s) during patches yielding 1.5µL water rewards (only trial-initial reward, top; four rewards, bottom) that were unstimulated (grey line, *n* = 1308 trials [top], *n* = 137 trials [bottom]) or coupled with dopamine (DA) stimulation (red dashed line, *n* = 1297 trials [top], *n* = 143 trials [bottom]). The corresponding average DA photometry signal is overlayed in cyan and orange. **B)** The local change in mouse speed (Δcm/s, top) and acceleration (Δcm/s^2^) during stimulated (red) vs. unstimulated (grey) 1.5µL water rewards on the first day and different day bins (left to right, mean ± s.d., two sample t-tests, *P* = 0.202 [speed D1], *P* = 0.66 [speed D1-6], *P* = 0.55 [speed D7-12], *P* = 0.443 [speed D13+], *P* = 0.201 [accel D1], *P* = 0.023 [accel D1-6], *P* = 0.895 [accel D7-12], *P* = 0.071 [D13+]).

**Figure S6.**
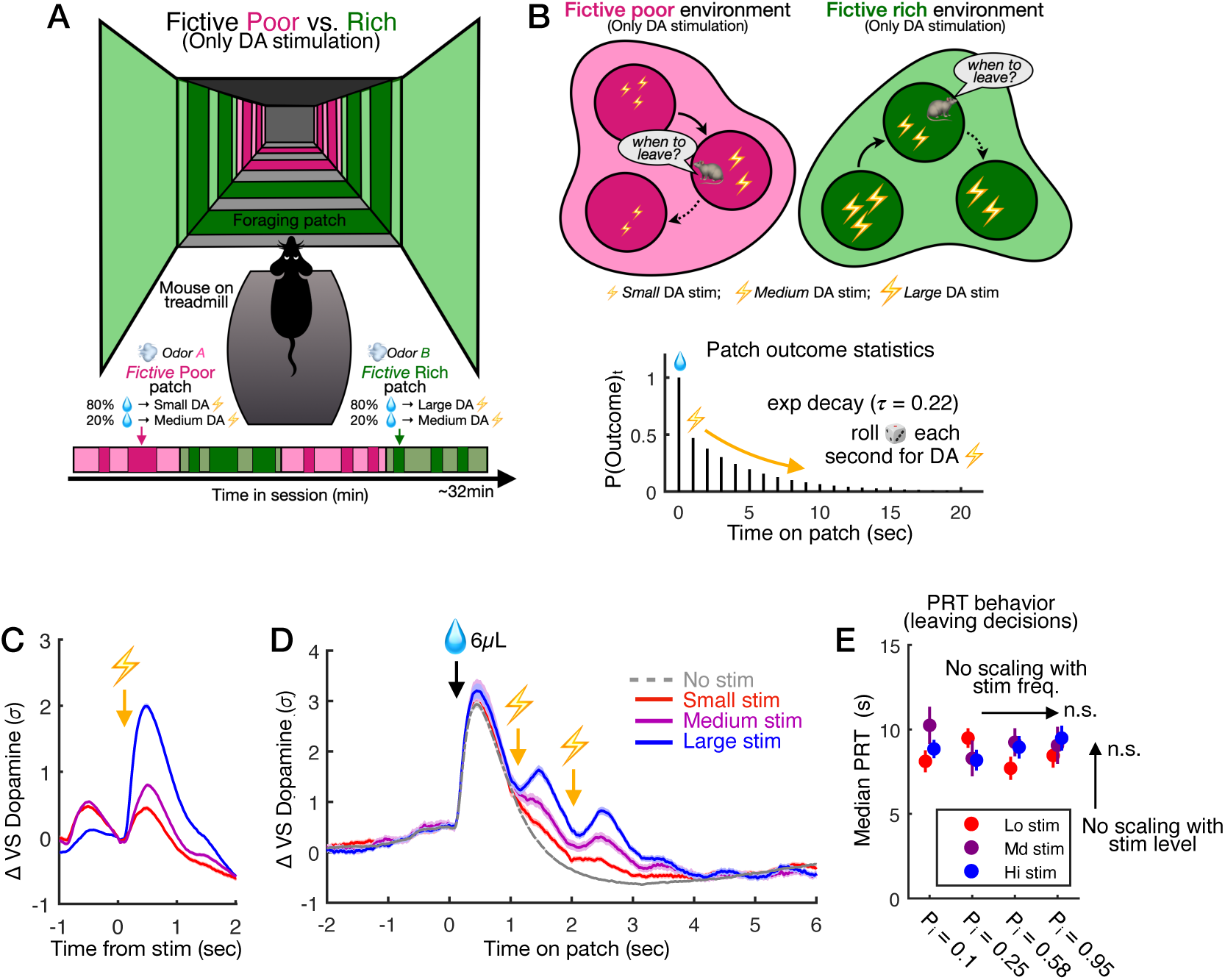
Immediate effects of phasic dopamine stimulation alone on patch leaving decisions. **A)** Design of fictive poor vs. rich environment experiment using phasic dopamine (DA) stimulation alone. Mice forage in two virtual environments which contain patches that all yield one trial-initial 6µL water reward followed by DA stimulation at three different levels to replicate the amplitudes of 1.5µL, 3µL, 6µL water responses (i.e. small, medium, and large phasic stimulation, creating fully fictive reward signals). **B)** The fictive poor (pink) environment contains patches yielding small or medium DA stimulation, and the fictive rich (green) environment contains patches yielding medium or large DA stimulation. The statistics of DA stimulation on foraging patches follows the same decaying probability structure as in the main natural reward task (Figure 1D). **C)** Ventral striatal (VS) DA signals aligned to stimulation alone at three different levels, producing reward-like transients that approximate responses to small (1.5µL, *n* = 3623 trials), medium (3µL, *n* = 3108 trials), and large (6µL, *n* = 2248 trials) water rewards (mean ± SEM). **D)** VS DA signals across the patch duration aligned to the trial-initial 6µL reward for small (red, *n* = 296 trials), medium (purple, *n* = 148 trials), or large (blue, *n* = 257 trials) patches in which the mouse received stimulation (i.e. fictive reward signals) at t = 1 and t = 2s, or no stimulation (grey, *n* = 1284 trials, mean ± SEM). **E)** Empirical PRTs measuring time invested on patches with increasing “fictive” reward (DA stimulation) frequency (initial reward probability at t = 1s, P_i_ on x-axis) and level (color), as in Figure 1F. Mice exhibit no scaling of patch leaving times (PRT) with more frequent or higher level stimulation (median ± s.d. for each size/probability patch type, multiple linear regression, *P* = 0.597 [reward size], *P* = 673 [reward probability], *n* = 3101 patch trials pooled across sessions).

**Table S1.** Comprehensive details for all statistical analysis.

| Figure | Comparison or association tested | Test(s) | Test Statistics | Effect size | 95% CI | P value | Observations/sample size | Significance label |
|---|---|---|---|---|---|---|---|---|
| Figure 1F | Reward size and freq, pooled PRT trials across mice/sessions, size effect | Multiple linear regression | β = 0.38471 | β = 0.38471 | [0.300, 0.469] | 4.82E-19 | d.f. = 22879 (trials) | *** |
| Figure 1F | Reward size and freq, pooled PRT trials across mice/sessions, freq effect | Multiple linear regression | β = 2.8084 | β = 2.8084 | [2.272, 3.345] | 1.28E-24 | d.f. = 22879 (trials) | *** |
| Figure 2A | PRT comparison poor vs. rich medium 3uL patches, early days (1-6) | Paired two sample t-test | t = -0.1259 | μ(Δ) = -0.0661 | [-1.1661, 1.0338] | 9.01E-01 | d.f. = 19 (n = 20 mice) | n.s. |
| Figure 2A | PRT comparison poor vs. rich medium 3uL patches, early days (7-12) | Paired two sample t-test | t = -0.1739 | μ(Δ) = -0.1272 | [-1.6578, 1.4034] | 0.8638 | d.f. = 19 (n = 20 mice) | n.s. |
| Figure 2A | PRT comparison poor vs. rich medium 3uL patches, early days (13-18) | Paired two sample t-test | t = -2.7658 | μ(Δ) = -1.1549 | [-2.0290, -0.2809] | 1.23E-02 | d.f. = 19 (n = 20 mice) | * |
| Figure 2B | PRT ratio rich/poor medium 3uL patches, early days (1-6) | One sample t-test | t = -0.5441 | μ = 0.9674 | [-0.1580, 0.0928] | 5.93E-01 | d.f. = 19 (n = 20 mice) | n.s. |
| Figure 2B | PRT ratio rich/poor medium 3uL patches, early days (7-12) | One sample t-test | t = 1.5399 | μ = 1.0848 | [-0.2366, 0.4062] | 0.5871 | d.f. = 19 (n = 20 mice) | n.s. |
| Figure 2B | PRT ratio rich/poor medium 3uL patches, early days (13-18) | One sample t-test | t = 2.2194 | μ = 1.3077 | [0.0175, 0.5979] | 3.88E-02 | d.f. = 19 (n = 20 mice) | * |
| Figure 2E | PRT regression performance, full models R2 | Wilcoxon signed-rank test | Z = 3.9199 | median(Δ) = 0.1990 | [0.1317, 0.2625] | 8.86E-05 | n = 20 mice | *** |
| Figure 2E | PRT regression performance, cv-R2 (10-fold) | Wilcoxon signed-rank test | Z = 3.9199 | median(Δ) = 0.1774 | [0.0967, 0.2456] | 8.86E-05 | n = 20 mice | *** |
| Figure 2F | PRT multiple regression, stats on mouse-level regressor coefs: Day | One sample t-test | t = 1.5312 | μ = 0.0671 | [-0.0246, 0.1588] | 1.42E-01 | d.f. = 19 (n = 20 mice) | n.s. |
| Figure 2F | PRT multiple regression, stats on mouse-level regressor coefs: Session | One sample t-test | t = 5.1792 | μ = 0.0803 | [0.0478, 0.1127] | 5.34E-05 | d.f. = 19 (n = 20 mice) | *** |
| Figure 2F | PRT multiple regression, stats on mouse-level regressor coefs: Env time | One sample t-test | t = 0.629 | μ = 0.008 | [-0.0187, 0.0348] | 5.37E-01 | d.f. = 19 (n = 20 mice) | n.s. |
| Figure 2F | PRT multiple regression, stats on mouse-level regressor coefs: Patch time | One sample t-test | t = -1.3455 | μ = -0.0214 | [-0.0548, 0.0119] | 0.1943 | d.f. = 19 (n = 20 mice) | n.s. |
| Figure 2F | PRT multiple regression, stats on mouse-level regressor coefs: Rrate global | One sample t-test | t = -2.7462 | μ = -0.0569 | [-0.1003, -0.0135] | 1.28E-02 | d.f. = 19 (n = 20 mice) | *** |
| Figure 2F | PRT multiple regression, stats on mouse-level regressor coefs: Rrate 10-back | One sample t-test | t = -6.2609 | μ = -0.06 | [-0.0801, -0.0399] | 5.18E-06 | d.f. = 19 (n = 20 mice) | *** |
| Figure 2F | PRT multiple regression, stats on mouse-level regressor coefs: Rrate 5-back | One sample t-test | t = -4.7469 | μ = -0.0555 | [-0.0800, -0.0310] | 1.40E-04 | d.f. = 19 (n = 20 mice) | *** |
| Figure 2F | PRT multiple regression, stats on mouse-level regressor coefs: Rrate 3-back | One sample t-test | t = -9.717 | μ = -0.0756 | [-0.0919, -0.0593] | 8.34E-09 | d.f. = 19 (n = 20 mice) | *** |
| Figure 2F | PRT multiple regression, stats on mouse-level regressor coefs: Rrate 2-back | One sample t-test | t = -9.4539 | μ = -0.0871 | [-0.1064, -0.0678] | 1.29E-08 | d.f. = 19 (n = 20 mice) | *** |
| Figure 2F | PRT multiple regression, stats on mouse-level regressor coefs: Rrate 1-back | One sample t-test | t = -7.9701 | μ = -0.1058 | [-0.1335, -0.0780] | 1.77E-07 | d.f. = 19 (n = 20 mice) | *** |
| Figure 2F | PRT multiple regression, stats on mouse-level regressor coefs: Rsize current | One sample t-test | t = 8.1322 | μ = 0.1311 | [0.0974, 0.1649] | 1.31E-07 | d.f. = 19 (n = 20 mice) | *** |
| Figure 2F | PRT multiple regression, stats on mouse-level regressor coefs: Rfreq current | One sample t-test | t = 2.8719 | μ = 0.0913 | [0.0248, 0.1579] | 0.0098 | d.f. = 19 (n = 20 mice) | ** |
| Figure 2F | PRT multiple regression, stats on mouse-level regressor coefs: Rich env | One sample t-test | t = 5.4727 | μ = 0.0975 | [0.0602, 0.1348] | 2.80E-05 | d.f. = 19 (n = 20 mice) | *** |
| Figure 2G | Rich environment effect as GLM regressor (converted to seconds) | One sample t-test | t = 4.4759 | μ = 0.8161; s = 0.8154 | [0.4345, 1.1978] | 2.59E-04 | d.f. = 19 (n = 20 mice) | *** |
| Figure 3E | Decision model comparison, kernel vs. flat | Two sample t-test | t = -2.1552 | Δ μ = -0.0476 | [-0.0938, -0.0014] | 0.0442 | d.f. = 19 (n = 20 mice) | * |
| Figure 3E | Decision model comparison, kernel vs. none | Two sample t-test | t = -6.2493 | Δ μ = -0.1529 | [-0.2041, -0.1017] | 5.31E-06 | d.f. = 19 (n = 20 mice) | *** |
| Figure 3G | Reward rate coef over time, early vs. late | Wilcoxon signed-rank test | Z = 2.3893 | median(Δ) = 8.7761 | [1.0142, 20.9698] | 0.0169 | d.f. = 19 (n = 20 mice) | * |
| Figure 3H | Compare decay taus for exponential across envs, early | Bootstrap shuffle env ID for time bins | n/a | Δ amp coef = 2.6248 | n/a | 0.052 | 1000 shuffles | n.s. |
| Figure 3H | Compare decay taus for exponential across envs, middle | Bootstrap shuffle env ID for time bins | n/a | Δ amp coef = 5.77 | n/a | 0.002 | 1000 shuffles | ** |
| Figure 3H | Compare decay taus for exponential across envs, late | Bootstrap shuffle env ID for time bins | n/a | Δ amp coef = 13.5050 | n/a | 0.002 | 1000 shuffles | ** |
| Figure 4E | Comparison of 3uL R1 response across poor vs. rich, late days (13-18) | Wilcoxon signed-rank test | W = 55 | median(Δ) = 0.8782 | [0.4503, 1.5624] | 0.002 | n = 10 mice | ** |
| Figure 4F | DA R1 differential timecourse (day bins) | Linear regression | R² = 0.313; β = -0.56624 | β = -0.56624 | [-0.810, -0.323] | 2.40E-05 | d.f. = 48 (50 measurements) | *** |
| Figure 4H | Comparison of 3uL omission across poor vs. rich, late days (13-18) | Wilcoxon signed-rank test | W = 44 | median(Δ) = 0.9241 | [0.4498, 1.1975] | 0.0078 | n = 10 mice | ** |
| Figure 4I | DA omission differential timecourse (day bins) (PRT > 3s) | Linear regression | R² = 0.158; β = -0.34618 | β = -0.34618 | [-0.620, -0.072] | 0.0147 | d.f. = 35 (37 measurements) | * |
| Figure 4K | Mouse level correlation between delta R1 DA and PRT bias | Linear regression | R² = 0.774; β = 5.4447 | β = 5.4447 | [3.043, 7.846] | 7.95E-04 | d.f. = 8 (10 mice) | *** |
| Figure 5E | PRT ratio FR/control medium 3uL patches, early days (1-6) | Two sample t-test | t = -0.8856 | Δ μ = -0.9641 | [-3.3898, 1.4615] | 3.97E-01 | d.f. = 10 (n = 11 mice) | n.s. |
| Figure 5E | PRT ratio FR/control medium 3uL patches, early days (7-12) | Two sample t-test | t = -1.1937 | Δ μ = -2.0625 | [-5.9126, 1.7875] | 0.2602 | d.f. = 10 (n = 11 mice) | n.s. |
| Figure 5E | PRT ratio FR/control medium 3uL patches, early days (13-18) | Two sample t-test | t = -2.2326 | Δ μ = -2.7392 | [-5.4730, -0.0054] | 4.96E-02 | d.f. = 10 (n = 11 mice) | * |
| Figure 5F | PRT ratio for 3uL patches in FR/Ctrl environment (chrimson mice) | One sample t-test | t = 3.0269 | μ = 0.2892 | [0.0763, 0.5020] | 1.27E-02 | d.f. = 10 (n = 11 mice) | * |
| Figure 5F | PRT ratio for 3uL patches in FR/Ctrl environment (tdTomato mice) | One sample t-test | t = 0.9230 | μ = 0.2056 | [-0.5033, 0.9144] | 0.4241 | d.f. = 3 (n = 4 mice) | n.s. |
| Figure | Comparison or association tested | Test(s) | Test Statistics | Effect size | 95% CI | P value | Observations/sample size | Significance label |
| Figure S1C | MVT simulated PRT - one threshold (3uL rich vs. poor) | Two sample t-test | t = 0.3218 | Δ μ = 0.0225 | [-0.1146, 0.1596] | 0.7476 | d.f. = 1998 (1000 trials each) | n.s. |
| Figure S1E | MVT simulated PRT - two thresholds (3uL rich vs. poor) | Two sample t-test | t = 5.6845 | Δ μ = 0.4012 | [0.2628, 0.5396] | 1.50E-08 | d.f. = 1998 (1000 trials each) | *** |
| Figure S2A | Scatter of PRT distribution | n/a | n/a | μ = 8.5382; s = 4.4862 | n/a | n/a | n = 20 mice | n/a |
| Figure S2B | Linear regression of mouse PRTs across even/odd day splits | Linear regression | R² = 0.774; β = 0.908 | β = 0.908 | [0.665, 1.151] | 3.14E-07 | d.f. = 18 (20 mice) | *** |
| Figure S2C | Scatter of engagement distribution | n/a | n/a | μ = 76.3822; s = 16.9956 | n/a | n/a | n = 20 mice | n/a |
| Figure S2D | Linear regression of patch engagement across even/odd day splits | Linear regression | R² = 0.856; β = 1.0986 | β = 1.0986 | [0.876, 1.322] | 5.21E-09 | d.f. = 18 (20 mice) | *** |
| Figure S2F | Comparison of patch trial engagement, rich vs. poor patches | Wilcoxon signed-rank test | Z = -3.8453 | median(Δ) = -0.1380 | [-0.2023, -0.0705] | 1.20E-04 | n = 20 mice | *** |
| Figure S2G | Mean rewards/trial for P3 Det vs. P3 Stc patches | Wilcoxon signed-rank test | Z = 2.9493 | median(Δ) = 3.2288 | [0.2947, 8.4417] | 3.20E-03 | n = 20 mice | *** |
| Figure S3C | Comparison of env entry DA across poor vs. rich, late days (13-18) | Wilcoxon signed-rank test | W = 3 | median(Δ) = -0.8640 | [-1.1143, 0.0474] | 0.0098 | n = 10 mice | ** |
| Figure S3D | Env entry differential timecourse (day bins) | Linear regression | R² = 0.165; β = 0.16813 | β = 0.16813 | [0.0558, 0.2805] | 0.00421 | d.f. = 46 (48 measurements) | ** |
| Figure S3G | Tau for inferred Vt from DA amplitudes (poor) | Exponential fit | n/a | tau = 0.266 | [0.2005, 0.3314] | n/a | n = 5 timepoint measures | n/a |
| Figure S3G | Tau for inferred Vt from DA amplitudes (rich) | Exponential fit | n/a | tau = 0.2625 | [0.1194, 0.4055] | n/a | n = 5 mtimepoint measures | n/a |
| Figure S5B | Delta speed comparison, stim vs. unstim rewards, day 1 | Two sample t-test | t = 1.2776 | Δ μ = 0.9657 | [-0.5190, 2.4504] | 0.2019 | d.f. = 561 | n.s. |
| Figure S5B | Delta speed comparison, stim vs. unstim rewards, days 1-6 | Two sample t-test | t = -0.4395 | Δ μ = -0.171 | [-0.9336, 0.5917] | 0.6603 | d.f. = 3771 | n.s. |
| Figure S5B | Delta speed comparison, stim vs. unstim rewards, days 7-12 | Two sample t-test | t = -0.6661 | Δ μ = -0.2711 | [-1.0690, 0.5268] | 5.05E-01 | d.f. = 4461 | n.s. |
| Figure S5B | Delta speed comparison, stim vs. unstim rewards, days 13-18 | Two sample t-test | t = -0.7664 | Δ μ = -0.3407 | [-1.2121, 0.5307] | 4.43E-01 | d.f. = 7442 | n.s. |
| Figure S5B | Delta accel comparison, stim vs. unstim rewards, day 1 | Two sample t-test | t = 1.2735 | Δ μ = 0.0333 | [-0.0181, 0.0847] | 2.03E-01 | d.f. = 561 | n.s. |
| Figure S5B | Delta accel comparison, stim vs. unstim rewards, days 1-6 | Two sample t-test | t = 2.2832 | Δ μ = 0.0239 | [0.0034, 0.0444] | 2.25E-02 | d.f. = 3771 | * |
| Figure S5B | Delta accel comparison, stim vs. unstim rewards, days 7-12 | Two sample t-test | t = -0.1320 | Δ μ = -0.0013 | [-0.0208, 0.0181] | 8.95E-01 | d.f. = 4461 | n.s. |
| Figure S5B | Delta accel comparison, stim vs. unstim rewards, days 13-18 | Two sample t-test | t = -1.8076 | Δ μ = -0.0166 | [-0.0345, 0.0014] | 7.07E-02 | d.f. = 7442 | n.s. |
| Figure S6E | Reward size and freq, pooled PRT trials across mice/sessions, size effect | Multiple linear regression | β = -0.0539 | β = -0.0539 | [-0.254, 0.146] | 5.97E-01 | d.f. = 3101 (trials) | n.s. |
| Figure S6E | Reward size and freq, pooled PRT trials across mice/sessions, freq effect | Multiple linear regression | β = 0.2726 | β = 0.2726 | [-0.994, 1.539] | 6.73E-01 | d.f. = 3101 (trials) | n.s. |

**Table S2.**
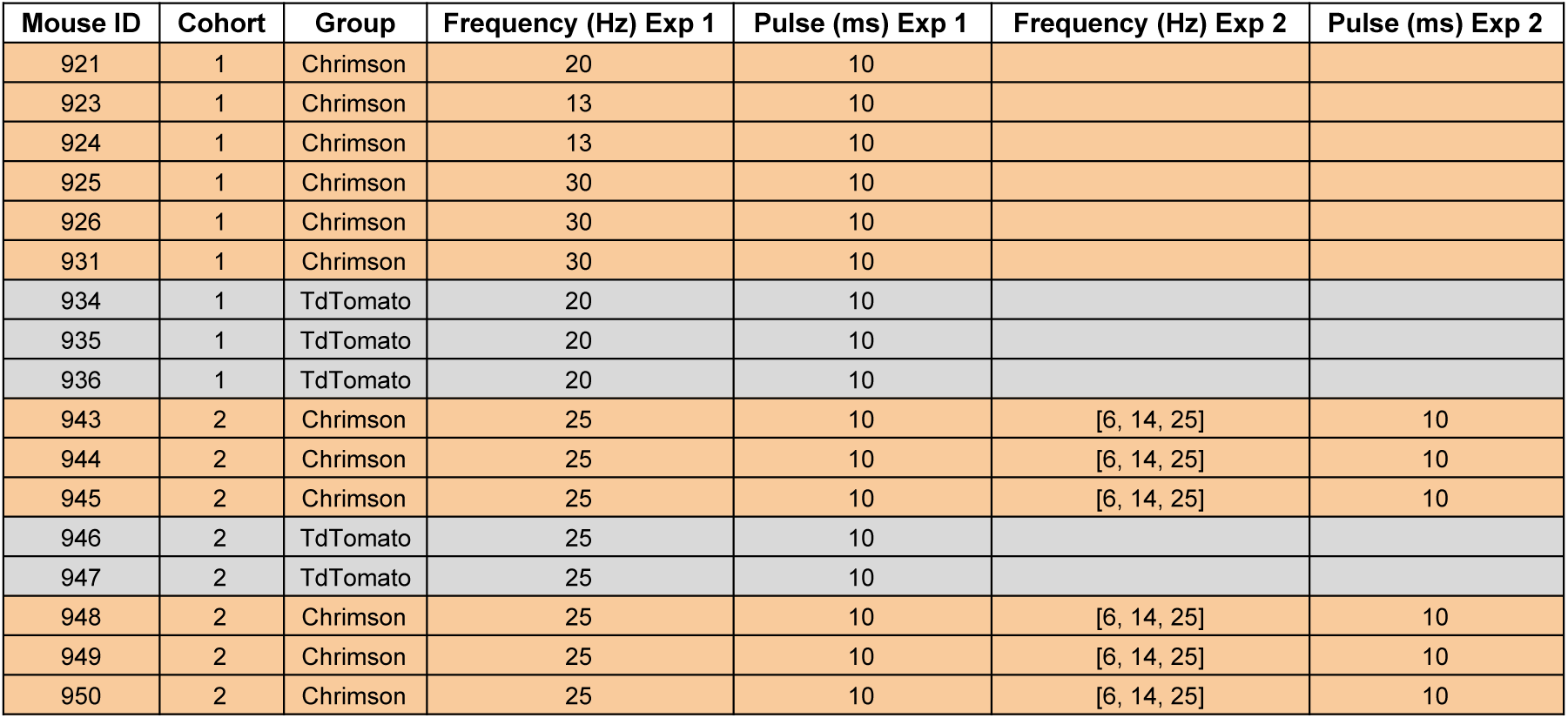
Summary of calibration parameters for dopamine neuron optical stimulation experiments.

| Mouse ID | Cohort | Group | Frequency (Hz) Exp 1 | Pulse (ms) Exp 1 | Frequency (Hz) Exp 2 | Pulse (ms) Exp 2 |
|---|---|---|---|---|---|---|
| 921 | 1 | Chrimson | 20 | 10 |  |  |
| 923 | 1 | Chrimson | 13 | 10 |  |  |
| 924 | 1 | Chrimson | 13 | 10 |  |  |
| 925 | 1 | Chrimson | 30 | 10 |  |  |
| 926 | 1 | Chrimson | 30 | 10 |  |  |
| 931 | 1 | Chrimson | 30 | 10 |  |  |
| 934 | 1 | TdTomato | 20 | 10 |  |  |
| 935 | 1 | TdTomato | 20 | 10 |  |  |
| 936 | 1 | TdTomato | 20 | 10 |  |  |
| 943 | 2 | Chrimson | 25 | 10 | [6, 14, 25] | 10 |
| 944 | 2 | Chrimson | 25 | 10 | [6, 14, 25] | 10 |
| 945 | 2 | Chrimson | 25 | 10 | [6, 14, 25] | 10 |
| 946 | 2 | TdTomato | 25 | 10 |  |  |
| 947 | 2 | TdTomato | 25 | 10 |  |  |
| 948 | 2 | Chrimson | 25 | 10 | [6, 14, 25] | 10 |
| 949 | 2 | Chrimson | 25 | 10 | [6, 14, 25] | 10 |
| 950 | 2 | Chrimson | 25 | 10 | [6, 14, 25] | 10 |

